# Proteomic and Genomic Signatures of Repeat-instability in Cancer and Adjacent Normal Tissues

**DOI:** 10.1101/491423

**Authors:** Erez Persi, Davide Prandi, Yuri I. Wolf, Yair Pozniak, Christopher Barbieri, Paola Gasperini, Himisha Beltran, Bishoy M. Faltas, Mark A. Rubin, Tamar Geiger, Eugene V. Koonin, Francesca Demichelis, David Horn

## Abstract

Repetitive sequences are hotspots of evolution at multiple levels. However, due to technical difficulties involved in their assembly and analysis, the role of repeats in tumor evolution is poorly understood. We developed a rigorous motif-based methodology to quantify variations in the repeat content of proteomes and genomes, directly from proteomic and genomic raw sequence data, and applied it to analyze a wide range of tumors and normal tissues. We identify high similarity between the repeat-instability in tumors and their patient-matched normal tissues, but also tumor-specific signatures, both in protein expression and in the genome, that strongly correlate with cancer progression and robustly predict the tumorigenic state. In a patient, the hierarchy of genomic repeat instability signatures accurately reconstructs tumor evolution, with primary tumors differentiated from metastases. We find an inverse relationship between repeat-instability and point mutation load, within and across patients, and independently of other somatic aberrations. Thus, repeat-instability is a distinct, transient and compensatory adaptive mechanism in tumor evolution.

## Introduction

Cancer clonal evolution (Cairns, 1975; Nowell, 1976) is marked by a wide range of genomic instabilities and somatic aberrations which lead to intratumor heterogeneity and eventually enable tumor cells to proliferate and metastasize (Stratton et al., 2009; Greaves & Maley, 2012; Yates & Campbell, 2012; Burrell et al., 2013). These aberrations include substantial, complex structural variation on every scale, as exemplified by the prevalence of aneuploidy (Gordon et al., 2012) and chromosomal instability (Lengauer et al., 1998; Bakum & Compton, 2012), hypermutation (Roberts & Gordenin, 2014; Campbell et al., 2017) and microsatellite instability (Wooster et al., 1994; Popat et al., 2005; Hause et al., 2016), complex short insertions and deletions (Ye et al., 2016), as well as large complex genomic rearrangements, such as chromothripsis (Stephens et al., 2011) and chromoplexy (Baca et al., 2013). Elucidating the relationship between different mutational classes is critical for inferring the exact clonal composition and phylogeny of tumors (Prandi et al. 2014; Beerenwinkel et al., 2015, Jiang et al., 2016), and subsequently, to determine how different aberrations affect clinical outcome (Birkbak et al., 2011; Andor et al., 2016; Hause et al., 2016) and which of these are involved in resistance to treatment and metastases formation (Naxerova & Jain, 2015; Beltran et al., 2016; Faltas et al., 2016).

Notwithstanding recent advances, identification of structural variations of short repeats in protein sequences remains elusive. This is the case because of the general difficulty to identify diverse types of repeats in sequences which vary in length, level of divergence and periodicity, and because of the relatively short length of reads obtained with next generation sequencing (NGS), which creates major difficulties for the current assembly techniques (Treangen & Salzberg, 2011; El-Metwally et al., 2013; Nagarajan & Pop, 2013), exacerbated by various causes of sequencing errors and DNA damage (Chen et al, 2017). Consequently, variations in the compositional order of proteins (Marcotte et al., 1999; Persi & Horn, 2013), a large class of mutations, which includes runs of amino-acids, short tandem repeats, interspersed repeats, repetitive domains, and more generally, over-representation of motifs in low complexity regions (hereafter, collectively denoted Repeats) has not been systematically characterized in cancer. To date, microsatellites, a relatively minor subclass of repetitive sequences, comprised of tandem repeats of 1-5bp units, represent the only well-studied case (Wooster et al., 1994; Popat et al., 2005; Hause et al., 2016; Campbell et al., 2017).

Repeats in proteins are hotspots of protein and species evolution that emerge through replication slippage and recombination (Levinson & Gutman 1987; Charlesworth et al., 1994; Paques et al., 1998). Repetitive domains are building blocks of key macromolecular complexes, (e.g., nuclear pores (Hoelz et al, 2011) and proteasomes (Pick et al., 2009)), and play essential roles in a variety of biological processes, notably, transcription regulation, protein-protein interaction and immunity, as exemplified by the enormous variety of Zinc-finger (Klug & Rhodes, 1987), Ankyrin (Mosavi et al., 2004), WD40 (Neer et al., 1994) and Leucine-rich (Bell et al., 2003) repeats in animal proteins. Variations in the number of repeat units have been associated with acquisition of new functions and rapid evolution of complex phenotypic traits in diverse life forms (Fondon & Garner, 2004; Verstrepen et al., 2005; Kashi & King 2006; Gemayel et al., 2010; Chavali et al., 2017). This fast evolution of repeats comes at the cost of promoting genetic diseases, in particular, cancer and neurodegeneration (Karlin et al., 2002; Gatchel & Zoghbi, 2005; La Spada & Taylor, 2010), where repeat dynamics (mostly, expansion but in some cases, contraction) often correlates with disease severity (Pearson et al., 2005; López Castel et al., 2010). Evolution of new repeats is markedly accelerated following duplication and is largely driven by positive selection, highlighting their potential role as disease drivers in somatic evolution (Persi et al., 2016).

In light of the importance of repeats in rapid evolutionary processes, coupled with their demonstrated involvement in human pathology, we hypothesized that repeat dynamics might play a more important role in tumor evolution than presently realized, especially, given that, in tumors, repeat generation is likely to be enhanced due to impaired DNA replication (Loeb et al., 1974; Tomasetti et al., 2017) and repair (Duval & Hamelin, 2002). To test this hypothesis, we generalized the quantification of repetitive motifs beyond microsatellites, and developed a rigorous methodology to systematically quantify variations in the repeat content (repeat-instability) of genomes (bypassing difficulties associated with assembly) and in the expression of repeat-containing proteins, directly from genomic and proteomic sequence raw data.

We applied the developed methodology to a collection of diverse datasets (Table 1), including (i) a proteomic dataset of breast cancer patients, (ii) genomic datasets of prostate cancer patients, including an original cohort of benign tissues, serving as a non-cancerous control, (iii) genomic pan-cancer cohorts from The Cancer Genome Atlas (TCGA) which include a tumor sample, an adjacent matched-normal control and a blood sample from each individual, providing for a comparison between tissues, and (iv) samples from patients with metastatic spread which allowed analysis of repeat-instability during the evolution from the primary tumor to the metastatic state. We demonstrate the utility of the methodology for identifying tissue-specific and tumor-specific repeat-instability signatures in cancer and normal tissues, and elucidate the dynamics and role of repeat-instability in tumor evolution.

**Table 1:** Datasets analyzed in this study. **1)** In the Proteomic dataset of 21 breast cancer patients [Pozniak et al, 2016], for each patient a matched-normal sample and a corresponding primary tumor (stage II or III) were measured. For 10 patients, with primary tumor stage III, a metastatic sample from the lymph nodes was also obtained. The 21 patients were obtained from two separate experimental pools of 10 and 11 patients each. Each pool contains patients from all stages (stage II, and stage III + metastases). **2)** In Barbieri dataset [Barbieri et al, 2012], for each of the 111 patients a tumor sample and a corresponding matched-normal sample, either blood or an adjacent tissue, were measured. **3)** Dataset of a non-cancerous benign prostate hyperplasia (BPH) tissue. **4-7)** pan-cancer TCGA datasets containing a primary tumor, an adjacent matched-normal and a blood sample from each patient. **8-9)** Datasets of two individuals with the largest metastatic spread, identified in two previous studies [Beltran et al, 2016; Faltas et al, 2016];

| Study/Tissue | No. of patients | Normal Samples |  | Cancer Samples |  | No. of Samples | Notes | Source / Ref |
| --- | --- | --- | --- | --- | --- | --- | --- | --- |
|  |  | Adjacent | Blood | Primary | Met |  |  |  |
| 1) Proteomic/Breast | 21 | 21 |  | 21 | 10 | 52 | Two experimental pools of 10 and 11 patients. Primary: stages II and III. Metastases: Lymph nodes. | Pozniak et al, 2016 |
| 2) WES/Prostate | 111 | 89 | 22 | 111 |  | 222 |  | Barbieri et al, 2012 |
| 3) WES/Prostate | 15 | 15 | 15 |  |  | 30 | Benign prostate hyperplasia (BPH) | Liu D, in preparation |
| 4) WES/Prostate | 41 | 41 | 41 | 41 |  | 123 | Trio | TCGA |
| 5) WES/Breast | 13 | 13 | 13 | 13 |  | 39 | Trio | TCGA |
| 6) WES/Bladder | 17 | 17 | 17 | 17 |  | 51 | Trio | TCGA |
| 7) WES/Lung | 42 | 42 | 42 | 42 |  | 126 | Trio | TCGA |
| 8) WES/prostate | 1 |  | 1 | 4 | 5 | 10 | Individual with metastatic spread. | Beltran et al, 2016 |
| 9) WES/Bladder | 1 |  | 1 | 5 | 7 | 13 | Individual with metastatic spread. | Faltas et al, 2016 |

## Results

### Measuring repeat-instability in amino-acid and nucleotide sequences

To measure the repetitiveness of motifs (*k*-mers) in a set of sequences, we define the compositional order ratio (CR) of a motif, as the total number of the motif recurrences divided by the total number of sequences in which the given motif appears (Figure 1a and Methods). The CR is high when a motif recurs multiple times in a sequence. Sequences in which motifs are highly recurrent are regarded as compositionally ordered. The CR signal tested on the human proteome strongly departs from the random expectation (Figure 1b), and furthermore, CR is substantially more robust for repeat identification than alternative measures of repetitiveness, such as the frequency of a motif or its fraction in the proteome (**Figure S1**).

**Figure 1:**
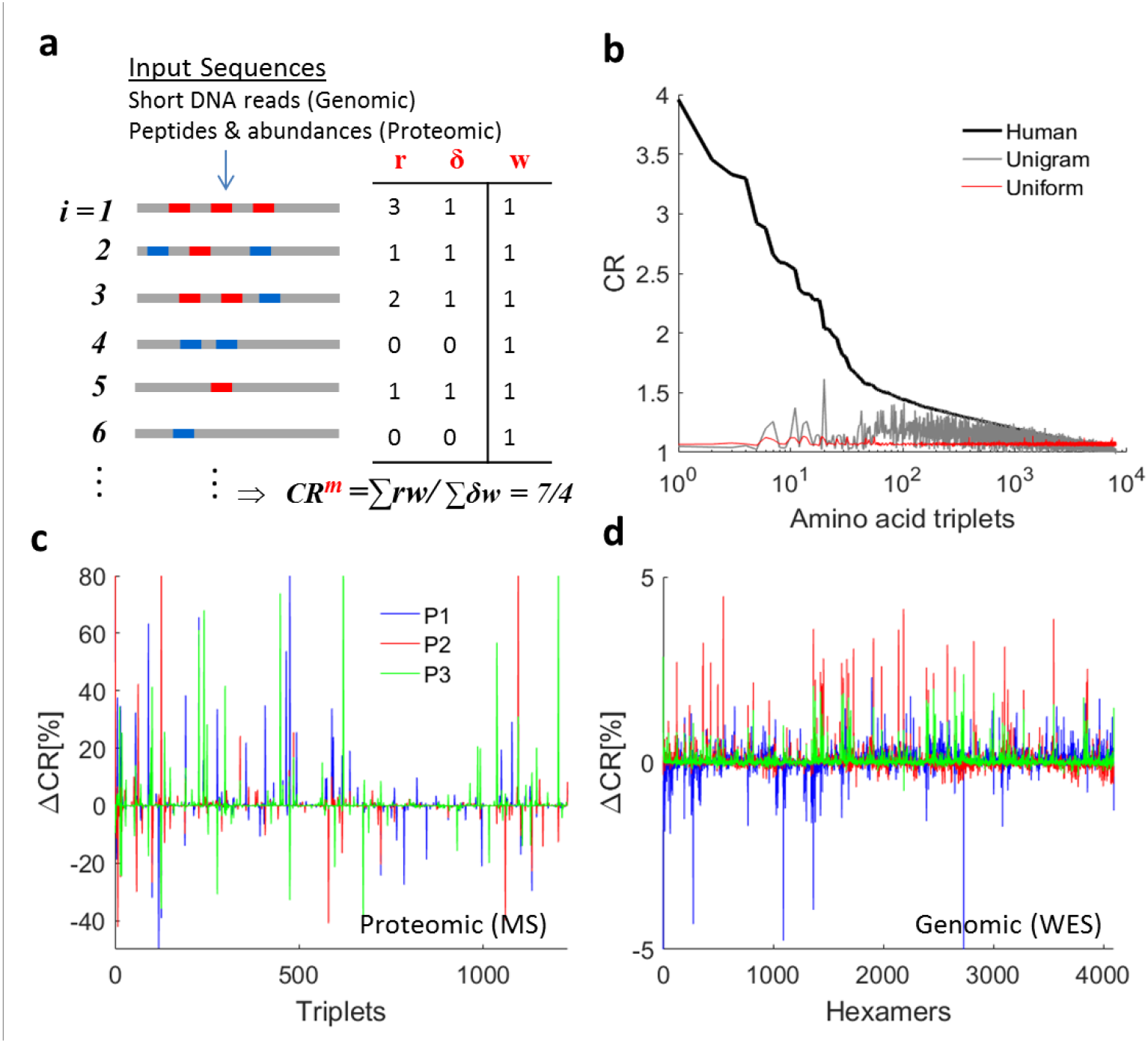
Methodology for estimating repeat-instability in genomic and proteomic sequence raw data. **A)** Illustration of the method for estimating the compositional order ratio (CR) of motifs (i.e., *k*-mers), using *k*=3 (triplets) for proteomic data and *k*=6 (hexamers) for genomic data (Methods). The distribution of two motifs (blue and red) is illustrated on a list of input sequences, *i=1.,.N.* Input sequences can be either peptides (with abundances *w_i_*) obtained from proteomic MS or DNA short-reads obtained from NGS (with equal abundances, *w*_*i*_=1). CR evaluation of the red motif in the case of genomic data is shown in the table, where for each sequence *i, r_i_* is the number of red motif recurrences, *δ_i_=1* if the red motif exists or *δ_i_=0* if no red motif exists. **B)** Application of CR estimation to the 8000 amino-acid triplet motifs distributed in the human proteome (black curve) shown against two random proteomes (identical protein length distribution), (i) uniform: with uniform probability of amino-acid recurrence (red) and (ii) unigram: where the probability of amino acids recurrence is based on the human proteome (grey). **C-D)** Examples of the *repeat instability* signatures (RIS=ΔCR) in tumors relative to their matched-normal tissue, of 3 patients (blue, red and green) in the breast cancer proteomic dataset (C) and in the genomic breast cancer dataset (D) (Table 1).

The CR can be directly estimated from both proteomic and genomic raw data (Figure 1a). In proteomic data, CR is evaluated from a list of ~100K measured peptides (typically 10-30 amino-acids long) and their abundances; the abundance values are used to estimate the effective number of sequences. We used triplets (*k*=3, 8000 amino-acid triplets) to measure CR in proteomic data, which is the optimal choice of motif length to characterize protein repeats (Persi & Horn, 2013). In whole exome sequencing (WES) data (coverage depth 100X), CR is computed from a list of ~100-200M short DNA reads (typically, 50-150 base-pairs long), using hexamers (*k*=6, 4096 nucleotide hexamers), such that the proteomic and genomic motif spaces have comparable sizes. The choice of *k*=6, a shorter unit length than the naïve choice of *k*=9 which translates into an amino-acid triplet, is also justified by the occurrence of synonymous substitutions that do not change the amino-acid composition (Methods). For CR evaluation, motifs can overlap and do not need to recur in tandem, such that all types of repeats, both pure and diverged, from runs to repetitive domains, that are shorter than half of the short-read length, can be identified (Methods). Several examples of repeats in proteins and their respective coding nucleotide repeats, identified in our analysis, are shown below, emphasizing the diversity of repeats that can be captured. Analysis of genomic data demonstrates that CR is a stable measure, which saturates at a low coverage depth (**Figure S2**) and is unbiased with respect to the sample size (**Figure S3**).

We define the **repeat-instability signature** (RIS=Δ**CR**) of a sample as the sum of the CR percentage changes for all motifs compared to a control sample (Methods). In a patient, the somatic signature of a tumor is computed relative to a control sample, taken either from an adjacent matched-normal tissue or from the blood. Figure 1c-d shows examples of typical tumor signatures in proteomic and genomic data. Because repeats can expand or contract in a given genome, we evaluate the **overall repeat-instability** by the sum over the absolute value of the signatures of all motifs (ORI=Ʃ|Δ**CR**|). Importantly, proteomic signatures reflect the compound effect of somatic genome instability and differential expression of repeat-containing peptides, whereas genomic signatures reflect genome instability alone. We applied this methodology to analyze peptide sequences in the proteomic dataset and short-read nucleotide sequences in the genomic (WES) datasets (Table 1).

### Proteomic repeat-instability reflects breast cancer tumor progression

We first applied the repeat analysis methodology to the proteomic dataset from 21 breast cancer patients (Pozniak et al., 2016) (Methods and Table1). We found that the CR of motifs (amino-acid triplets) tends to increase in tumors relative to matched-normal tissues, as measured by the average signature of triplets across patients (Figure 2a). To ensure that this trend was not a consequence of large variations of CR in a few patients, we assessed the frequency of variation in the CR among the patients. The histogram of the frequencies is bimodal, with CR consistently increasing for many triplets and consistently decreasing for a few triplets among the patients (Figure 2b). The remarkable shift to high frequency that was observed for strongly altered (>1%) triplets confirms that CR-increase is the dominant phenomenon and CR can be used to characterize tumors. Principal component analysis (PCA) of the CR estimates (52 samples *x* 1229 triplets) shows clear separation of matched-normal samples from tumor samples in the first 2 principal components (Figure 2c). This separation was captured in two experimental pools (Table 1 and Methods), indicating that the tumor *vs* normal segregation is robust. The PCA analysis also suggests that the dimensionality of discrimination is low, such that classifiers can be built using a small number of discriminative features (i.e., triplets). Examples of discriminative triplets (e.g., `PVP`, `APV`, `APA`, `YGY`, `DVL`, `TAA`) are shown in **Figure S4**.

**Figure 2:**
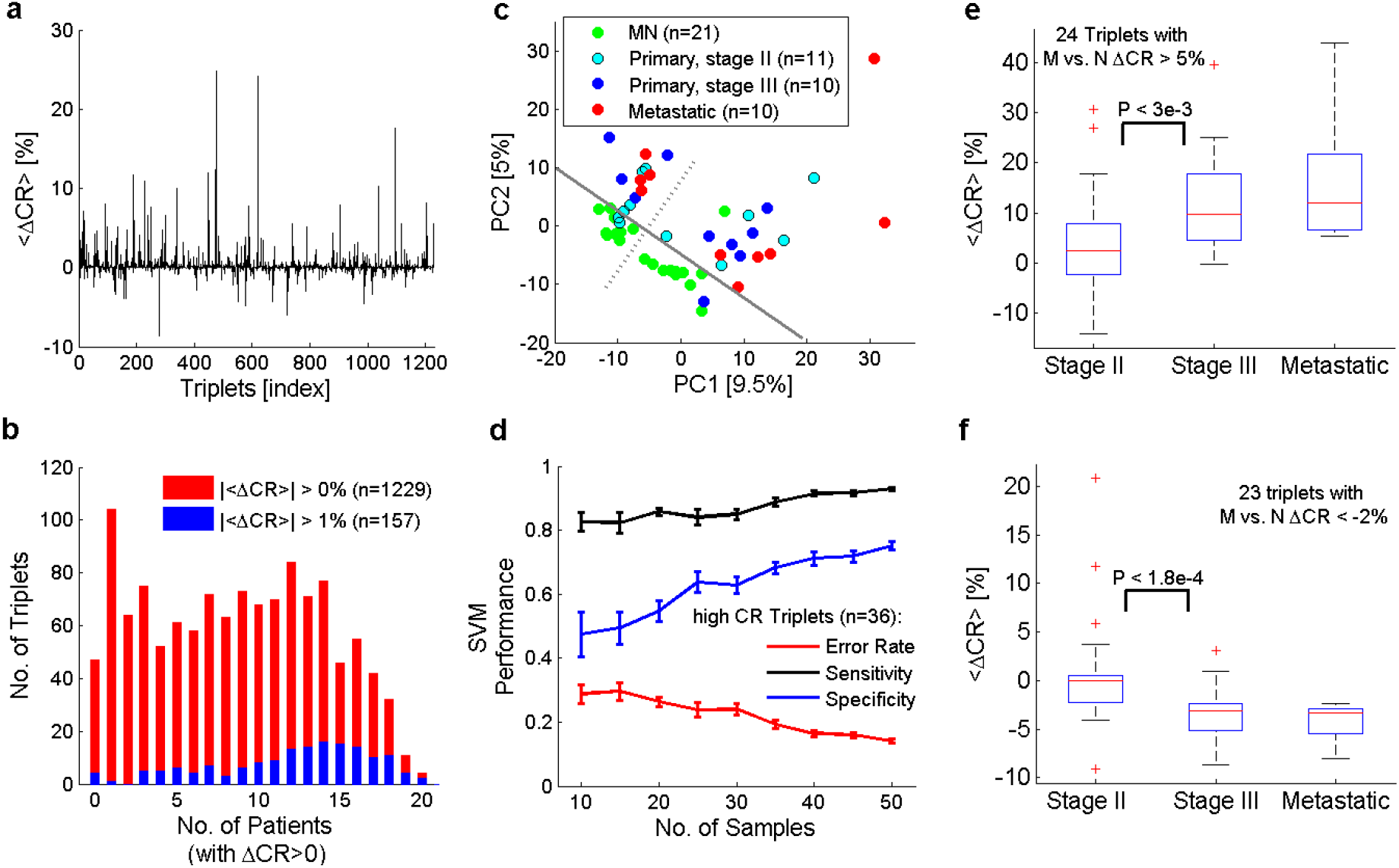
Analysis of tumor signatures in proteomic data from 21 breast cancer patients. **A)** Average repeat-instability in tumors, evaluated relative to the respective matched-normal (MN) tissue in each patient, demonstrates an overall tendency for CR increase. When two tumor tissues are sampled from the same patient (i.e., stage III and metastatic), the average signature is used. **B)** Histograms of the number of triplets vs. the number of patients in which the triplets’ CR increased (ΔCR>0), shown for all identified triplets (|<ΔCR>| > 0%, n=1229, red) and for triplets with |<ΔCR>| > 1% (blue, n=157). Frequencies of positive variation of triplets are the x-axis values divided by 21. **C)** PCA analysis of the CR matrix (52 samples *x* 1229 triplets), indicating the separability between normal and tumor samples in the first two principal components. A grey solid line is superimposed for visual clarity of the discrimination. Perpendicular to it, the dashed grey line indicates the division between the two experimental pools (see Methods and Table 1). Seperability between tumor and normal tissues is robust. **D)** The effect of sample size on the SVM linear classifier, using the top high-CR triplets as features (n=36). Classification performance in a leave-one-out analysis improves with sample size. Error bars are estimated from 20 trials of random choice of samples at each point on the *x*-axis. **E)** Triplets (n=24) with increased average signature (<ΔCR> > 5%) in the metastatic samples (M) relative to matched-normal (N) reveals that CR increases in the transition from stage II to stage III. **F)** Triplets with decreased signatures in the metastatic signature (<ΔCR> < −2%, n=23) tend to decrease from stage II to stage III. P-values correspond to the Kolmogorov-Smirnov test. Trends in (E) and (F) are not expected at random (**Figure S5**). Accuracy = percentage of correct classifications. Sensitivity = TP/(TP+FN). Specificity = TN/(TN+ FP). TP = true positives, TN = true negatives, FP = false positives, FN = false negatives.

To further test the predictive signal of repeat-instability signatures, we built binary classifiers that discriminate between normal and tumor samples, using support vector machine (SVM) with a linear kernel, and examined various feature-selection criteria in a standard leave-one-out analysis (Methods, **Table S1**). Every tested selection criterion (Kolmogorov-Smirnov test, Fisher-score and CR-based criteria) achieved classification accuracy >80% with a small set of triplets (~10-30). We further inspected the simplest criterion for selecting triplets with high CR, i.e. those that frequently recur in the list of identified peptides. This approach achieved a maximum accuracy of 89% with only 36 selected triplets (**Table S1**) that are frequently and significantly altered among the patients (**Figure S4**). To ensure that the classifier performance is not sensitive to the small number of samples, we also tested its performance as a function of the number of samples, and found that it improves as more samples are included, testifying to the generality and robustness of this simple approach for discriminating between tumor and normal samples (Figure 2d).

Although good performance was achieved in discriminating tumor from normal samples, metastases do not appear to be well separated from primary tumors (Figure 2c). Nonetheless, we noticed that several triplets displayed consistent variation from normal to primary tumor to metastases (e.g., the triplet `TAA` in **Figure S4**). Thus, to test for signatures correlated with cancer progression, we selected triplets with the strongest average signature in the metastases (relative to matched-normal), and tested whether their signatures varied from stage II to stage III. This particular comparison was performed because selection of triplets with strong signatures in the metastases statistically selects weaker signatures in stages II and III, but differences between stage II and stage III are not expected to be affected (**Figure S5**). As implied by the tendency of CR-increase in tumors, we found more triplets with average signatures increased from stage II to stage III (Figure 2e) than triplets for which the average signatures decreased (Figure 2f). These trends were robust to the choice of the threshold used to select triplets with strong signatures in metastases (**Figure S5**). Notably, a weaker variation between stage III and metastases was observed, suggesting that the differential expression of repeats is especially important at early stages of tumor evolution. Mapping all discriminative triplets to proteins and identifying repeat-unstable proteins (Methods) indicated that the proteins with high repeat-instability are enriched among the proteins encoded by known cancer genes (**Figure S6**) which is compatible with a role of repeat instability in oncogenesis.

The proteomic CR signatures reflect changes in expression levels of repeat-containing proteins and, accordingly, do not directly convey any information on genomic somatic variation in the repeat content in protein-coding DNA. Hence, to explore the role of repeat-instability in somatic evolution, we turn to the analysis of genomic data. Hereafter, we analyze short reads of nucleotide sequences obtained from WES data (Table 1).

### Genomic repeat-instability discriminates between healthy and cancerous prostate tissues

We explored genomic repeat-instability in prostates, the tissue type with the richest dataset among the ones examined, which includes samples from both healthy individuals and cancer patients (Table1). We analyzed the TCGA dataset of prostate cancer patients (Cancer Genome Atlas Research Network, 2015), focusing on cases for which a primary tumor sample and two control samples, from blood and from an adjacent matched-normal tissue, were collected. In each patient (n=41), we computed the CR signatures of the tumor and of the adjacent normal samples relative to blood. In most of the patients, the signatures of tumor and adjacent normal samples were closely similar, displaying strong positive correlation (Figure 3a and **Figure S7**). This correlation was independent of the shape of the signatures (i.e., expansion-dominated or contraction-dominated), implying that the signatures are primarily tissue-specific (**Figure S7**). Furthermore, high similarity between tumors and normal samples is also observed across patients, as demonstrated by the bimodal distribution of the pairwise correlations (**Figure S7**), which reflects the prevalence of positive or negative correlation between the signatures from different patients, a consequence of the dominance of either repeat expansion or contraction, in a given genome.

**Figure 3:**
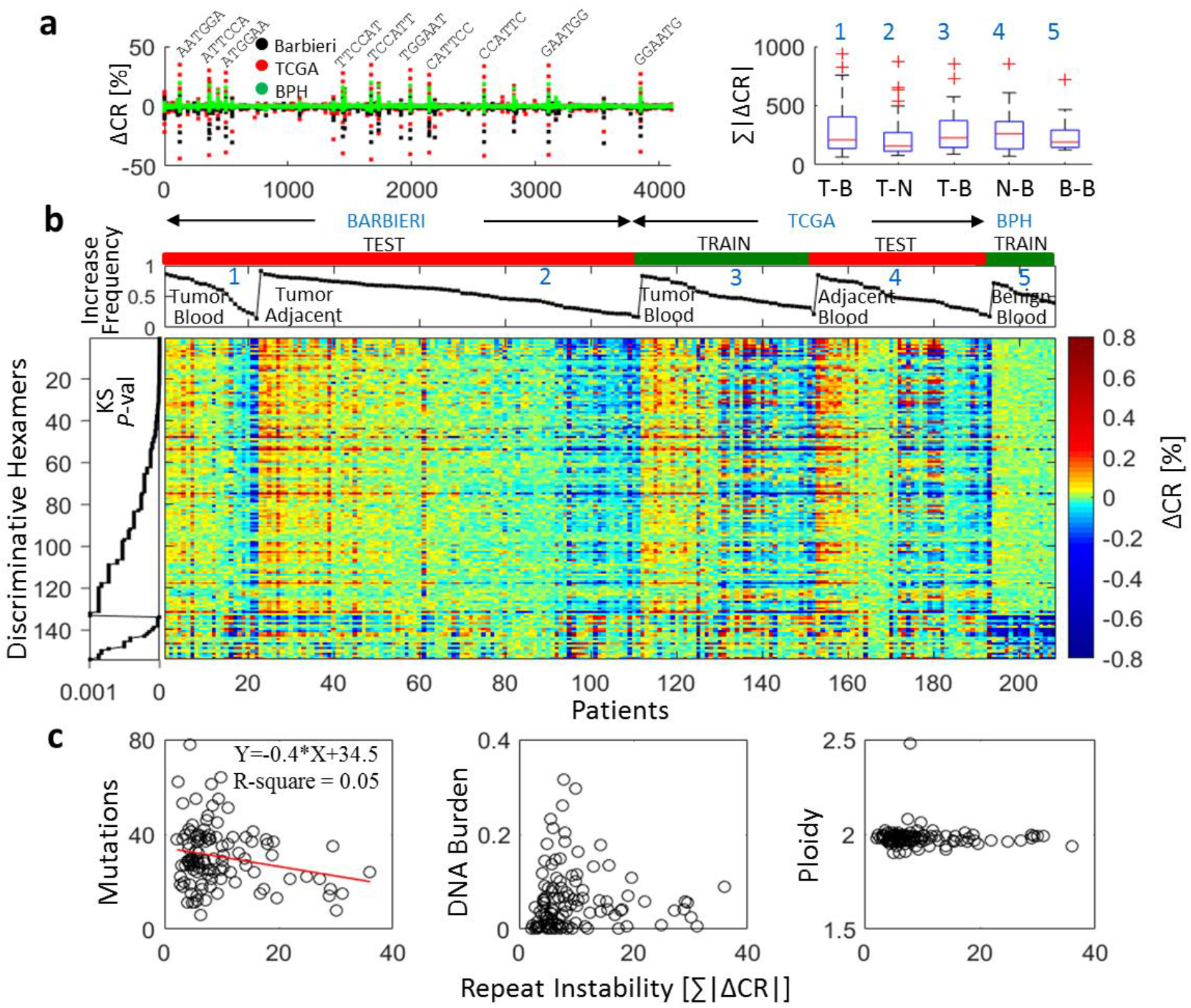
Repeat-instability in prostate tissues. **A)** Repeat-instability signatures (RIS=ΔCR) of all prostate datasets: Barbieri (n=111, black), TCGA (n=41, red) and Benign prostate hyperplasia (BPH, n=15, green) superimposed (*Left*). 10 most dominant hexamers are shown. The overall repeat instabilities (ORI=Ʃ|ΔCR|) distributions of tumors (T) and adjacent normal tissues (N), in the different prostate datasets (*Right*). In the Barbieri dataset, tumor signatures are estimated relative to blood (T-B, n=22) or to an adjacent tissue (T-N, n=89), respectively (1-2). In TCGA, signatures of tumors (T-B) and adjacent normal tissues (N-B) are computed relative to blood (3-4). BPH signature (B-B) is computed relative to blood (5). **B)** Heat map of the 154 discriminative hexamers found in Table 2 (task 1) comparing RIS of tumors in the TCGA dataset with that of benign prostates, computed relative to the blood (Train). Test sets display similar characteristics to tumors of the train set (signatures ID as in A, *right*). Results of training-test sets are robust (**Figure S8**). Colors reflect ΔCR values. Hexamers are ordered by their KS *P*-values, grouped into those that have higher |ΔCR| in tumors and those with higher |ΔCR| in the benign tissues within the train set. Patients are ordered by the portion of discriminative hexamers that increased in each signature. **C)** Relationship between ORI and different genomic variables in tumors. Each point represents a patient in the Barbieri dataset. ORI is estimated using the set of 154 discriminative hexamers.

**Table 2:** Results of linear binary SVM classifiers, applied to the genomic prostate datasets. For each discrimination task (1-3) a standard leave-one-out test was performed. **A-B)** Datasets and the corresponding compared signatures (|ΔCR|) of two groups of patients. **C-E)** SVM performance without feature selection (i.e., considering all 4096 hexamers). **F-H)** The number of discriminative hexamers found by Kolmogorov-Smirnov test comparing the distribution of |ΔCR| between the two groups in each task using as selection criterion *P*-value < 0.001. Hexamers are separated into those whose |ΔCR| distribution is higher in the first group (i.e., the tumor signature) and those whose |ΔCR| is smaller in the first group. 62 out of the 96 hexamers in task 3 overlap the 154 hexamers identified in task 1. Accuracy = percentage of correct classifications; Sensitivity = TP/(TP+FN); Specificity = TN/(TN+ FP), where TP = true positives, TN = true negatives, FP = false positives, FN = false negatives. See **Figure S9** for the predictive signal of the discriminative hexamers.

| Task | Comparison<br>Group1 vs. Group2 | | Linear SVM Classifier<br>(Leave One Out) | | | No. of Discriminative motifs<br>(KS $P$-value < 0.001) | | |
| --- | --- | --- | --- | --- | --- | --- | --- | --- |
| | A)<br>Datasets | B)<br>\|ΔCR\| Signatures | C)<br>Accuracy | D)<br>Sensitivity | E)<br>Specificity | F)<br>Larger in<br>Group 1 | G)<br>Smaller in<br>Group 1 | H)<br>Tot |
| 1 | TCGA<br>BPH | Tumor-Blood vs.<br>Benign-Blood | 98% | 1 | 0.93 | 133 | 21 | 154 |
| 2 | TCGA<br>TCGA | Tumor-Blood vs.<br>Adjacent-Blood | 34% | 0.31 | 0.38 | 0 | 0 | 0 |
| 3 | TCGA<br>BPH | Tumor-Adjacent<br>Benign-Blood | 98% | 1 | 0.93 | 79 | 17 | 96 |

To determine whether these repeat-instability signatures include tumor-specific characteristics, we compared them to signatures of benign prostate hyperplasia (BPH) from healthy individuals not affected by prostate cancer (n=15, here referred to as healthy individuals) that were computed relative to matched blood (Table 1). Superposition of tumor and benign signatures (Figure 3a, left panel) highlights strong similarity between prostate cancer and healthy tissues but the healthy signatures are weaker. This difference is recapitulated by the overall repeat-instability (ORI=Ʃ|ΔCR|), which shows high similarity between tumor and adjacent normal tissues in cancer patients, but a lower instability of the benign prostates in healthy individuals (Figure 3a, right panel). However, these differences had limited statistical significance, emphasizing the strong tissue specificity of the signatures. Thus, to identify tumor-specific features, we trained SVM classifiers as in the proteomic case but, to account for the expansion-dominated and contraction-dominated genomic signatures; we considered the absolute value (|ΔCR|) in our analysis. Tumor tissues were robustly discriminated from benign ones (Table 2, task 1), and 154 discriminative motifs were identified using Kolmogorov-Smirnov test (*P*-value < 0.001): 133 showed flat signatures and 21 exhibited strong contraction in benign tissues relative to tumors. The classifiers cannot distinguish between tumor and adjacent normal signatures when each is computed relative to blood, and no discriminative motifs were found (Table 2, task 2), as expected from their close similarity. Using adjacent normal tissues as a control in place of blood leads to significantly weaker signatures (Figure 3a). Nonetheless, these signatures contain tumor-specific information as the identified discriminative motifs between these signatures and those of benign tissues largely overlap those that were identified in task 1 (Table 2, task 3). Therefore, this comparison yields an attenuated tumor signature. We used this signature when a blood sample was missing.

To test the predictive power of the 154 discriminative motifs, we considered an independent dataset (Barbieri et al., 2012) of 111 prostate cancer patients (Table 1). The repeat-instabilities of the test and training sets showed remarkable similarity (Figure 3b). The similarity between tumor and adjacent matched-normal signatures, but not with healthy signatures, implies that the adjacent normal prostate tissues in cancer patients contain tumor-specific features, in the absence of histological evidence. We validated this prediction using various training-test sets, demonstrating that tumors are predicted with high accuracy (>90%), based on both tumor and adjacent normal signatures (**Figure S8**).

To explore which genes were most affected by repeat-instability, we mapped the short reads encompassing repeats from the Barbieri dataset onto the human genome, and estimated repeat-instability at the gene level (Methods). We found that the 10 most unstable motifs, with |ΔCR|>5% in >80% of the patients (Figures 3a), were not discriminative and did not map to coding regions, but rather to regulatory regions (**Figure S9**). Because these dominant signatures appear in all tissues, including benign prostates, it seems likely that they represent repeat hotspots in non-coding regulatory regions that might exert currently unknown effects on transcription regulation (Vinces et al., 2009). In contrast, discriminative motifs mapped to protein-coding regions (**Figure S9**). As in the proteomic case, the set of most repeat-unstable genes was significantly enriched in known cancer genes (**Figure S10**). We analyzed in detail the amino-acid and nucleotide compositions of the identified repeat-unstable genes. This analysis also emphasized the ability of our methodology to identify diverse types of repeats (Methods), from runs of amino-acids in proteins, as exemplified by the glutamine tracks in FOXP2 protein (**Figure S11**), to repetitive domains, as in the case of the Cysteine-rich PAK1 inhibitor CRIPAK (**Figure S12**), based on the recurrence of hexamers in the protein-coding DNA.

Lastly, we explored the relationship between repeat-instability and other somatic aberrations by querying the Barbieri dataset. The overall repeat-instability of discriminative motifs was weakly inversely related to the non-silent point mutation load, but was independent of copy-number alterations (i.e., DNA burden) and aneuploidy status (Figure 3c). The apparent, even if weak, tradeoff between repeat-instability and the mutation load suggests that, in tumor evolution, repeat-instability could be a compensatory mechanism for point mutations. To test this hypothesis, we performed a pan-cancer analysis, exploring a wider distribution of mutation loads.

### Genomic repeat-instability is inversely related to somatic point mutation load in the pan-cancer dataset

In addition to the prostate adenocarcinoma analyzed above, 3 additional cancer types were selected from TCGA (Table 1), such that the selected cancers represent different point mutation load regimes: prostate and breast cancers have relatively low numbers of point mutations per sample, and bladder and lung cancer have comparatively high numbers of point mutations (Lawrence et al., 2013). As in prostate cancer, we focused our analysis on patients with available data from all three types of samples (tumor tissue, adjacent matched-normal tissue, blood), to measure the repeat-instability signatures of the primary tumor and its adjacent normal sample relative to the blood sample. Similarly to the observation on prostate cancer (**Figure S7**), normal and tumor signatures were highly correlated in individual patients across all cancer types (**Figure S13**).

Across cancer types, we identified a consistent inverse relationship between the overall repeat-instability and the number of non-silent point mutations, which was independent of the overall genomic burden (Figure 4a). Despite the similarity between adjacent normal and tumor signatures observed in each patient (**Figure S13**), patients with low mutational load cancers (i.e., breast) display higher repeat-instability in normal tissues when compared to the respective tumor signatures, whereas for high mutational load cancers (bladder and lung), this trend is reversed. In prostate patients, the similarity between tumors and the normal tissues was the highest, explaining the difficulty in identifying tumor-specific features in this case, and the need for a non-cancerous control (cf. Figure 3). Thus, repeat instability is high when the mutation load is low and low when the mutation load is high, and this effect is stronger for the signatures of adjacent normal tissues in the vicinity of tumors than for the tumor signatures themselves. The inverse relationship between the point mutation load and the overall repeat instability holds both for gain (expansion) and loss (contraction) of repeats, but is more pronounced for gain (**Figure S14**).

**Figure 4:**
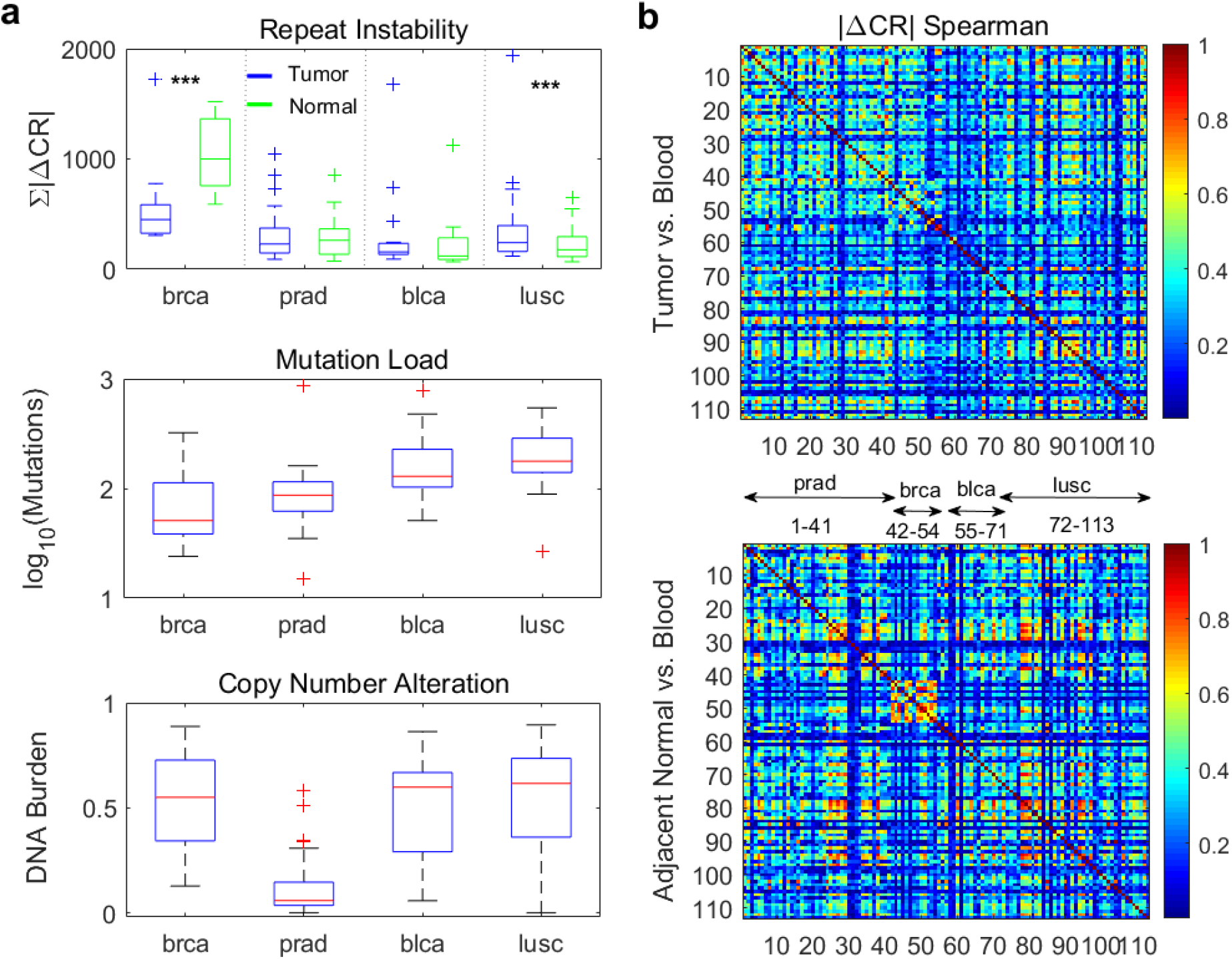
Pan-cancer analysis of repeat-instability in the TCGA datasets. **A)** The overall repeat-instability (ORI=Ʃ|ΔCR|, vs. blood) in tumors and corresponding adjacent matched-normal tissues (*top;* few outliers with ORI>2000 are omitted for clarity. See **Figure S14**). The mutation load, estimated by number of non-silent point mutations (*middle*) and the copy-number alterations (i.e., DNA burden) measured by the fraction of altered genes (gain or loss) in the proteome (*bottom*). An inverse relationship exists between repeat-instability and the point mutation load. In low mutational load cancer types repeat instability is larger in the adjacent normal tissues than in tumors, but this reverses in high mutational load cancers (*** indicates KS-test *P*-value < 0.01). **A)** Spearman correlation among patients of tumor signatures (*top*) and of adjacent matched-normal signatures (*bottom*) measured relative to the blood sample in each patient.

Further, to elucidate the differences between cancer and adjacent normal genomes across tissues, we assessed the pairwise correlations among patients of both adjacent normal signatures and of the respective tumor signatures (Figure 4b). Adjacent normal tissue signatures display higher correlations across patients compared with the respective tumor signatures, that is, the normal signatures are more tissue-specific. This effect was more pronounced in patients with low mutational load cancers (breast and prostate) where the repeat-instability is high. The weaker tissue specificity of tumor signatures suggests that common mechanisms, such as impaired DNA replication and repair, similarly affect the repeat content of tumors across tissues and individuals, and consequently, blur the similarity between the same tissue samples across individuals, with a net effect of reduced tissue-specificity. This effect conversely enhances the tumor-specific signal in tumor signatures, as we demonstrated in the case of prostate cancer (Figure 3 and Table 2), leading to a more homogenous structure of the pairwise correlations between tumor signatures, across patients and across different cancer types (Figure 4b).

### Genomic repeat-instability recapitulates tumor phylogeny within patients and correlates with metastatic spread

To substantiate the role of repeat-instability as a distinct and compensatory mutation class in tumor evolution, we studied two patients with metastatic spread (Table 1). The two patients with the largest number of available sequenced samples from different anatomical sites were selected from two recent studies of metastatic prostate cancer (Beltran et al., 2016) (WCM0) and chemotherapy-resistant urothelial carcinoma (Faltas et al., 2016) (WCM117), respectively. The prostate cancer patient represents a case of a low mutation load cancer type, whereas the bladder cancer patient represents a case of high mutation load cancer type.

The analysis of the repeat-instability signatures from different anatomical sites of the same patient, measured relative to blood, highlights a clear hierarchy based on the correlation between the signatures, where primary tumors and metastases are well separated into two clusters, both in the prostate cancer patient (Figure 5a) and in the bladder cancer patient (Figure 5b). Further, we approximated the tumor phylogeny by similarity dendrograms, inferred from the repeat-instability pairwise correlation distance, and from the Hamming distances between mutated genes, using the simple unweighted pair group method with arithmetic mean (UPGMA) to link samples. The repeat-instability and point mutation distance-based dendrograms were closely similar (Figure 5), and were weakly sensitive to the choice of either linkage (e.g., shortest distance or UPGMA) or distance methods (i.e., Hamming or Euclidian in the case of mutations, and Spearman or Pearson correlation in the case of repeat instability) (**Figure S15**).

**Figure 5:**
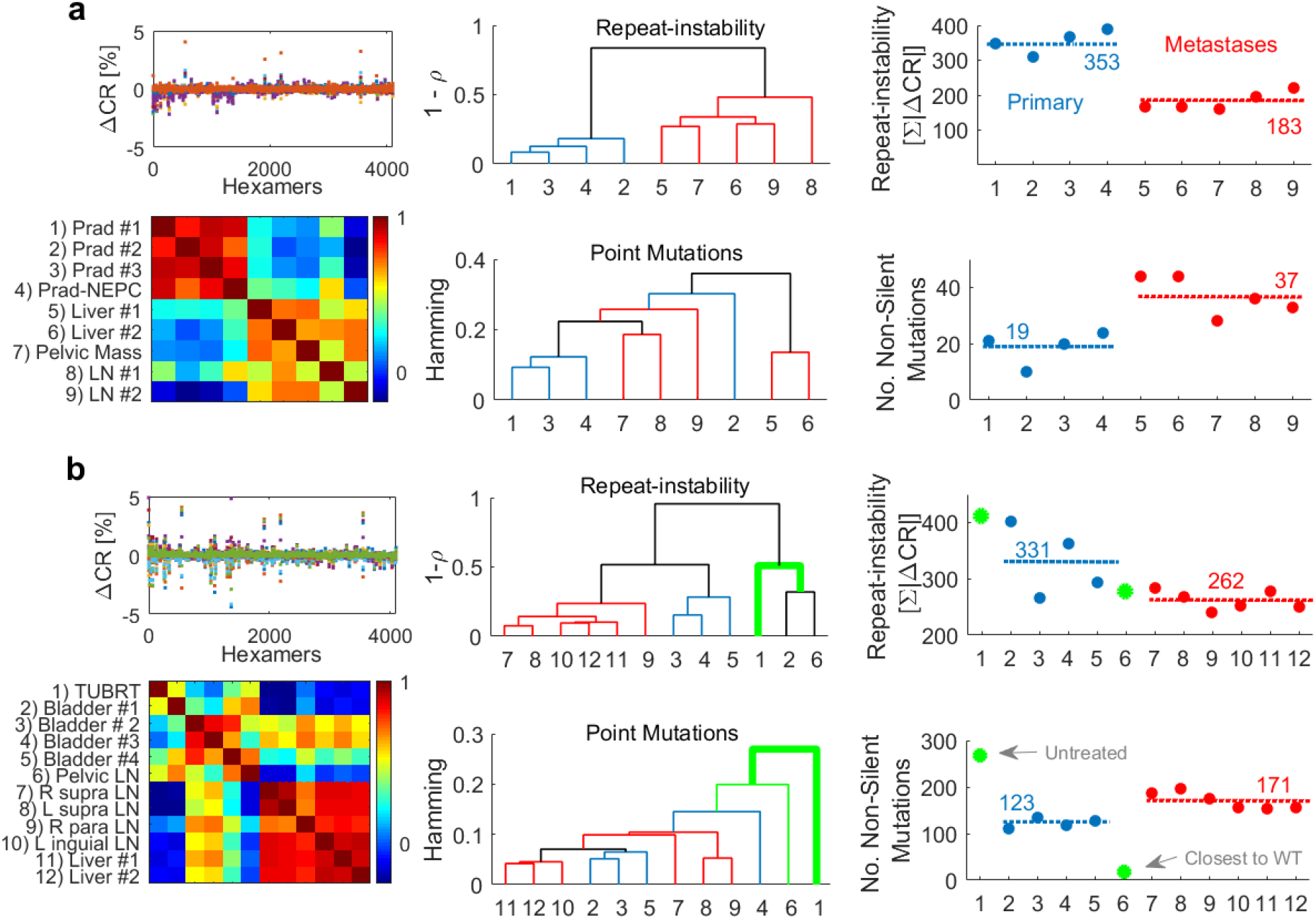
Analysis of multiple samples from two individuals with metastatic spread. **A)** Analysis of a prostate cancer patient, with the large metastatic spread and multiple available biopsies from [Beltran et al., 2016]. **Left panel**, the repeat-instabilities signatures (RIS=ΔCR) of different samples vs. blood is superimposed (top), and the heat map of the pairwise spearman correlations across samples (*bottom*). **Middle panel**, the dendrogram inferred by the correlations distance (1-ρ) among the RIS of samples (top), and the dendrogram inferred by the hamming distance between the non-silent point mutation of samples across genes (*bottom*). Dendrograms are estimated using unweighted pair group method with arithmetic mean (UPGMA). Primary tumor leafs are colored in blue, metastatic leafs in red, and connecting branches in black. **Right panel** depicts the inverse relationship between the overall repeat-instability (ORI=Ʃ|ΔCR|) (*top*) and the non-silent point mutation load (*bottom*). Averages values are depicted by dashed lines. **B)** Similar analysis of a bladder cancer patient, with the largest metastatic spread, following treatment, identified in [Faltas et al., 2016]. Untreated sample (#1) and a treated metastatic sample (Pelvic) which is the closest to the tumor ancestor wild-type (#6) are colored in green, and are not considered to estimate averages of repeat-instability and mutation load (dashed lines, *right*) of the treated samples.

The tumor phylogeny inferred from repeat-instability was concordant with the detailed phylogeny that we have previously obtained by rigorous analysis of the clonal composition of samples and the tempo of somatic aberrations (Beltran et al., 2016; Faltas et al., 2016). Repeat-instability based phylogeny captures some fine details of the relationship among samples: **(i)** the prostate primary tumor sample with neuroendocrine features (Figure 5a) is close to the other primary tumor samples (Beltran et al., 2016), **(ii)** the metastatic pelvic lymph node in the bladder cancer patient that was surgically removed at the time of the cystectomy, is close to the bladder primary tumors (Figure 5b), in particular, to the untreated primary tumor (despite marked difference in the overall repeat-instability between these samples), as previously inferred using independent techniques (Faltas et al., 2016) and, **(iii)** metastases from the same anatomical site cluster together, with few differences. The finding that the repeat-instability approximated phylogenies are closely similar to the true phylogenies suggests that repeat-instability evolves by divergence, at clock-like rates.

To test the hypothesis that repeat-instability is a compensatory adaptive path of tumor evolution, we compared the overall repeat-instability and the non-silent point mutation load in primary tumors and metastatic sites. In both patients, repeat-instability was higher in primary tumors than in metastases, corroborating the inverse relationship between repeat-instability and the point mutation load (Figure 5, right panels). These findings imply that repeat-instability is most pronounced at early stages of tumor progression, likely, acting as a transient genome alteration that compensates for the relatively low number of driver mutations in primary tumors, and is partially reversed as mutations accumulate and tumor cells adapt.

## Discussion

The involvement of repeat-instability in human pathology is supported by ample evidence (Wooster et al., 1994; Karlin et al., 2002; Gatchel & Zoghbi, 2005; Popat et al., 2005; Pearson et al., 2005; La Spada & Taylor, 2010; López Castel et al., 2010; Hause et al., 2016; Campbell et al., 2017, Chavali 2017). However, the current understanding of the role of this phenomenon in somatic evolution is mostly limited to microsatellites. In this study, we describe a method that measures the repeat content in the genome and in the proteome, directly from proteomic peptide data and genomic short-reads. Our approach accounts for a wide range of repeats. Applying this approach to an array of studies allowed us to assess repeat-instability across many patients and in different tissue types, comparing tumors with normal tissues, yielding insights into its role in tumor evolution.

Our analyses of genomic signatures show that, compared to blood, cancer and adjacent normal tissues often evolve similarly and manifest comparable repeat-instability signatures. Furthermore, such signatures are, to a large extent, tissue-specific, in accordance with previous studies of repeat-instability in other disorders (Pearson et al., 2005; López Castel et al., 2010). However, we also identified significant tumor-specific signatures, which correlate with the course of tumor evolution and allow for discriminating healthy samples from cancers. Specifically, we found that repeat-instability is inversely related to the point mutation load but is independent of aneuploidy and genomic burden of somatic gene copy number. This inverse relationship was observed between low mutational load cancers (prostate and breast) and high mutational load cancers (bladder and lung) primary tumors, and between primary tumors and metastases from the same patient. Given that repeat-instability includes microsatellites, our findings support and generalize the results of recent studies showing that microsatellite instability is prevalent across cancer types (Hause et al., 2016), but is consistently more pronounced in patients with low mutation loads compared to those with high mutation loads (Campbell et al., 2017).

Given that about two-third of the mutations in cancer can be attributed to replication errors (Tomasetti et al., 2017), which promote repeat instability (Pearson et al., 2005; López Castel et al., 2010, Campbell et al., 2017) the observed tissue specificity of repeat-instability signatures could be explained, at least, in part, by the tissue-specific cell division rates. Because blood has a relatively high rate of cell divisions (Tomasetti C, & Vogelstein, 2015), it is substantially diverged from other tissues, and is therefore an adequate choice as an outgroup control for characterizing repeat-instability (and other somatic aberrations). Also, the effective population size of blood cells is likely large, so that purifying selection is highly efficient. Therefore, blood is likely to be largely free of deleterious mutations, and hence, an adequate control. Conversely, the relatively low rates of cell divisions in bladder and lung tissues likely contribute to the overall lower repeat-instability in these tissues.

Collectively, these observations indicate that repeat-instability is a distinct adaptive path in tumor evolution. We propose a model of tumor evolution (Figure 6a) in which, at the initial phase of tumorigenesis (low mutational load cancers at the pan-cancer level, and primary tumors at the patient level), the number of cancer driver mutations is low, and repeat-instability is likely to serve as an additional, complementary mechanism, which increases (or maintains) the fitness of tumors. Later in tumor evolution, when metastases and/or high mutation load tumors accumulate more mutations and subsequently more drivers, repeat-instability is reduced as tumors adapt to their specific ecological niches. Specifically in high mutational load cancers, along with the accumulation of drivers, the number of deleterious passenger mutations substantially increases, imposing selective pressure (Bozic et al., 2010; McFarland et al., 2013) that could reduce repeat-instability. This theoretically predicted transition in evolutionary regime at high mutation load is also captured by the association of the point mutation load with clinical outcome (Persi et al., 2018). Thus, although most tumors evolve near neutrality (Williams et al., 2016; Weghorn & Sunyaev, 2017; Martincorena et al., 2017; Persi et al., 2018), high mutational load (specifically during the transition to a metastatic state) leads to decreased fitness, both through intracellular mechanisms and through generation of neo-antigens which elicit immune response (Berraondo et al., 2016; Li et al., 2016; Mlecnik et al., 2016a; Yarchoan et al., 2017a). The immune system, then, exercises purifying selection, thereby reducing repeat-instability in the tumor cell population. Those repeats that have been fixed in the cell population are likely beneficial, consistent with recent observation of microsatellite-unstable colon carcinomas, where strong purifying selection eliminates antigen-presenting tumors from the population (Mlecnik et al., 2016a), whereas immune-adapted tumors metastasize (Mlecnik et al., 2016b). Hence, although the accumulation of diverse types of mutations represents vulnerability for cancer, eventually, mutations that confer selective advantage are fixed in a population of tumor cells whereas deleterious mutations are removed, such that cancer maintains its fitness. The observed sharp decrease in repeat-instability appears to be a consequence of the dynamic nature of repeat propagation, being fast and reversible, unlike accumulation of point mutations. Although a bad omen, this evolutionary race also opens new avenues for identifying neo-antigens and developing immunotherapies against immune-adapted tumors (Berraondo et al., 2016; Yarchoan et al., 2017b).

**Figure 6:**
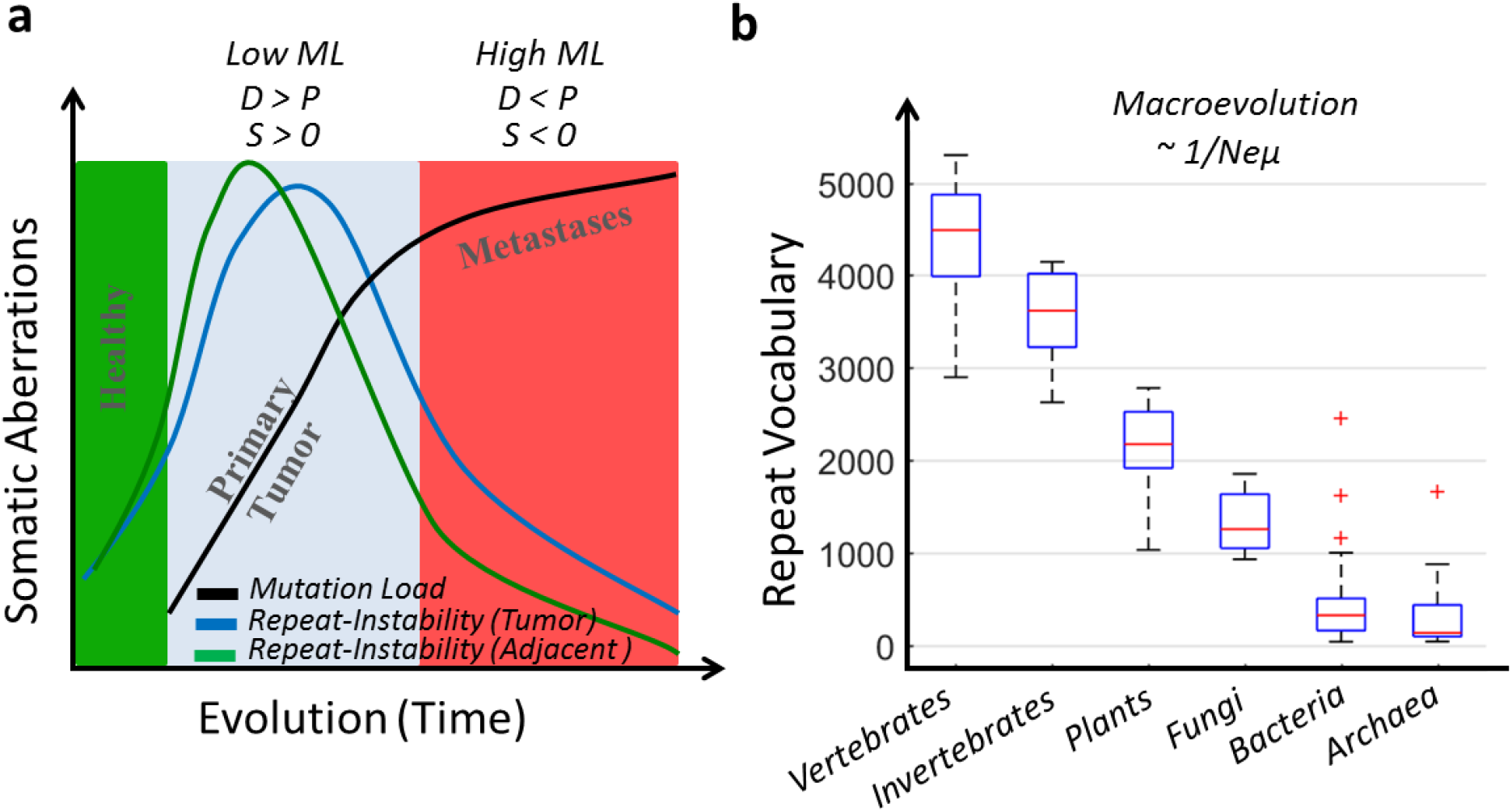
Proposed Evolutionary model of repeat dynamics in cancer and normal tissues. **A)** In healthy tissues (e.g., benign) repeat-instability is low. At the initial phase of tumor evolution (e.g., primary and low ML cancer types), tumors harbor a small number of positively selected (*S>0*) drivers (*D*). Repeat instability acts to increase or maintain the fitness of tumors. Tissues in the vicinity of tumors (e.g., adjacent presumed normal tissues) react similarly to the selective pressures imposed by the microenvironment (and therapy, if applied). Because they lack mutations, repeat instability is even larger than that of the corresponding tumors (that already have adapted), but is also quickly reduced as the transition to a neoplastic state is not achieved and normal cellular function is retained. Later in evolution (i.e., metastases and high ML cancer types), the number of driver mutations increases, tumors are more adapted and repeat instability reduces. At least in high mutation load cancers, the accumulation of passenger mutations (*P*) outcompete the drivers (*P>D);* hence, cancers resort to purifying selection (*S*<0), which reduces the repeat-instability. Hence, repeat-instability acts as a transient, compensatory mechanism. The faster transient effect in adjacent normal tissues explains their higher repeat-instability in low mutation load cancers and their lower repeat-instability in high mutation load cancers, relative to the respective tumors. **B)** Repeats content, measured as the vocabulary of amino-acid triplets that compose protein repeats, correlates with ordering of organisms by the product of effective population size and mutation rate (adapted from [Persi & Horn, 2013]).

According to our model, at the initial phase of tumor evolution, repeat-instability can compensate for the lack of sufficient number of cancer drivers and thus increase the tumor fitness, whereas later in evolution, high repeat-instability negatively affects tumor fitness and is selected against. A positive correlation between high microsatellite instability with better prognosis in cancer patients has been reported (Hause et al., 2016). In the context of our model, these findings are likely to reflect stages of tumor progression at which repeat-instability already exceeded the optimal value. The existence of compensatory adaptive paths, that is, point mutations vs. repeat-instability, suggests that, although the dynamic range of somatic aberrations in cancers is substantial, the fitness of tumors tends to be more stable over time and can be robust to environmental pressure.

Under this view of tumor evolution, adjacent normal tissues, residing in physical proximity to tumors, evolve under comparable selective pressures imposed by the microenvironment and therapy, so that they acquire tumor-specific signatures. Accordingly, analysis of these tissue samples should allow prediction of cancer breakout, prior to pathological evidence, as we demonstrated in the case of prostate cancer. This view is concordant with recent reports on significant similarities between the somatic signatures of cancers and normal tissues (Cooper et al., 2015, Martincorena et al., 2015). We hypothesize that repeat instability is low in healthy tissues, rapidly increases in tumors and adjacent normal tissues, and then is reduced as cancer progresses. This transient dynamics is partially captured by the single patient analysis, showing a large increase in the repeat-instability in the untreated primary tumor relative to WT, followed by a gradual reduction in the treated primary and metastatic tumors, and coupled to the increased point mutations (cf. Figure 5b). Such a compensatory transient mechanism of repeats in tumors is reminiscent of chromosomal duplications in Fungi (Yona et al., 2012) and gene duplications in viruses (Cone et al., 2017), which appear to represent the first, rapid route of adaptation. Adjacent normal tissues seem to exhibit an even faster dynamics of transient repeat instability than tumors. Conceivably, the cells in these tissues start on the path of tumorigenesis, but fail to undergo neoplastic transformation, whereby repeat-instability is rapidly reduced (Figure 6a). This explains the differences between repeat instabilities of adjacent normal and of tumors across tissues, showing a faster transient-like effect in adjacent normal tissues as function of the mutation load across cancer types (cf. Figure 4). This dynamics, leading to the link between repeat instability and cancer progression, is also concordant with the somatic evolution of repeat instability in many neurological disorders (Pearson et al., 2005; López Castel et al., 2010). Taking into account also the observed connection between proteomic repeat-instability and cancer progression, we suggest that repeat instability signatures can serve as important diagnostic and prognostic markers that could be sensitive enough to detect cancer in early stages.

In this work, we quantified and emphasized the importance of gain and loss of repeat units in tumor evolution. Similar to gene duplications (Lynch & Conery, 2000; Kondrashov et al., 2002), selective constraints are relaxed in new repeats following duplication (Persi et al., 2016), such that mutations can accumulate at higher rates and eventually lead to the acquisition of new functions. However, in contrast to gene duplicates, which evolve under only slightly relaxed purifying selection and mostly exhibit subfunctionalization of ancestral proteins (Kondrashov et al., 2002, Innan & Kondrashov, 2010), new repeats evolve much faster, under strongly relaxed purifying selection and positive selection, such that neofunctionalization is likely to be the primary route of evolution (Persi et al., 2016). Such rapid evolution of new repeats has been documented in colonic carcinogenesis (Ionov et al, 1993). Indeed, we observed that highly repeat-unstable genes were enriched among known cancer genes, both in genomic and proteomic data. This implies involvement of repeat-instability and, more specifically, fast-evolving new copies of repeats, in oncogenesis.

The microevolutionary dynamics of repeat instability in cancer (Figure 6a) is consistent with the evolution of repeats over long spans of evolution (Figure 6b). In diverse life forms, following the rapid evolution of new repeat copies (Persi et al., 2016), some repeats become conserved as they gain function (Schaper et al., 2014; Persi et al., 2016). The conservation of mutated repeats appears to eventually translate into an increase in the diversity of the repeat content of extant species proteomes, in a manner that correlates with the ordering of major clades by *N_e_* x *μ* (effective population size, *N_e_*, multiplied by the mutation rate, *μ*), that is, by the power of purifying selection (Lynch and Conery, 2003; Persi et al., 2013) (Figure 6b). These parallels with findings on species evolution should inform the study of repeat instability in somatic evolution of cancer.

Further integrated genomic-proteomic research is needed to study how somatic changes in the genomic DNA are translated into differential expression of repeat-containing peptides and how new copies diverge by accumulating mutations during tumor evolution. Such research could lead to the identification of new cancer drivers and the development of therapeutic strategies, in particular immunotherapies, which target this mutational class. Furthermore, given that our results indicate that repeat-instability is an adaptive mechanism that is important at the early stages of tumor evolution, we hypothesize that repeat instability signatures may be relevant for early cancer detection by cell-free DNA and liquid biopsy analysis as well as other methods. The role of repeat instability in other pathologies and evolutionary scenarios has yet to be explored.

## Methods

### Measuring Repeat Instability in Genomes and Proteomes

The extent of repetitiveness of a motif *m*, in a genome or proteome, is measured by its compositional order ratio (CR), as illustrated in Figure 1A, defined as the number of the motif recurrences in a set of sequences divided by the number of sequences in which it appears:

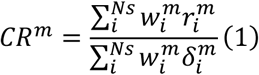

where *m* = 1, …, *N_m_*, and *N_m_* = *A^k^* is the number of searched *k*-long motifs over the alphabet *A* (e.g., *N_m_*=20^3^ for amino-acid triplets, *N_m_*=4^6^ for DNA hexamers). 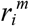 is the number of recurrences of motif *m* on sequence *i*. 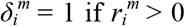 and is zero otherwise. 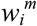 is a weight factor which measures the relative abundance of sequence *i* in a sample, particularly relevant for proteomic data described below. *N_s_* is the number of sequences in the data. By definition CR is ≥ 1.

The existence of compositionally ordered sequences in a proteome is responsible for the fact that some motifs will tend to recur multiple times in a relatively small number of sequences, and will therefore have high CR. For proteomic data, we use amino-acid triplets (*k*=3, *N_m_*=8000) which is the optimal motif length for characterizing protein repeats (Persi & Horn, 2013). The significance of CR evaluation is demonstrated in **Figure S1**, where we compare the SwissProt human reference proteome to random models (with identical length distribution of proteins), and compared CR with other measures such as the frequency of motifs or their fraction in the proteome (i.e., the number of sequences in which motifs appear).

The variation in the repetitiveness of a motif *m* between two samples, in particular, between a tumor tissue and its matched-normal tissue or a blood sample, is measured in percentages by:

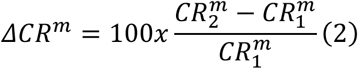

where 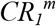 and 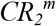 are the CR of motif *m* in samples 1 and 2, respectively. The repeat-instability signature is then expressed as the spectrum of ΔCR of all motifs, and the overall repeat-instability is given by the sum over the absolute values of motifs’ variations, Ʃ_m_|ΔCR_m_|.

To find an adequate motif length (i.e., *k*) for the analysis of genomic data, we first performed a systematic search of all DNA *k*-mers (with 3 ≤ *k* ≤ 9) on few patients. Variations in CR of motifs in DNA were not evident at *k*=3, whereas for large *k* (*k*>6), the vocabulary size is too large to allow a useful analysis. Hence, working with a motif vocabulary size comparable to that used in proteomic data, choosing *k*=6, such that *N_m_*=4096, was sufficient for the identification and characterization of repeat content at the DNA level. The choice of *k*=6 represents a shorter unit length than the naïve choice of *k*=9, which translates into an amino-acid triplet. As nucleotide nonamers can harbor many synonymous substitutions that do not change the amino-acid composition, even a tandem array composed of exact copies of amino-acid triplets is not expected to be coded by exact copies of nucleotide nonamers. The lengths of recurring motifs at the DNA level are thus expected to be shorter than 9. Conversely, repetitive hexamers in the protein-coding DNA may identify repeats that are not observed in protein sequences, due to insertions and deletions.

To validate that the identification of repetitive hexamers in nucleotide sequences significantly overlap the repeats in proteins (identified by repetitive triplets) we analyzed the repeat content of all human proteins. We employed the compositional order methodology (Persi & Horn, 2013) to both the amino-acid sequences using triplets and to the corresponding protein-coding DNA using hexamers. In random sequences, the probability of identifying a motif that recurs more than *n* time in a sequence of length *L* is determined by the Bernoulli distribution: 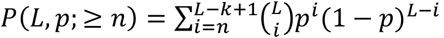, where *p* is the probability of selecting a *k*-mer over an alphabet *A*, such that *p*=*1*/*A^k^* (i.e., 1/8000 for amino-acids and 1/4096 for nucleotides). Hence, for a given *k* and *A, n* can be set to ensure a comparable statistical significance of identifying a recurrent element. By this consideration, we searched the human proteome for amino-acid triplets that recur at least 5 times in a protein, and nucleotide hexamers that recur at least 8 times in the respective protein-coding DNA. This analysis largely identified the same (~5000) repeat-containing proteins (Jaccard score 0.6). Hence, although there are obvious differences between the results of searching for repetitive triplets in amino-acid sequences and searching for repetitive hexamers in corresponding nucleotide sequences, basing the large-scale analysis in this study on k=3 and k=6, respectively, appears as a reasonable choice.

Evaluation of the number of motif recurrences in sequences allows overlap, such that all types of repeats can be identified. For example, in the sequence <u>AAAAAA</u>AAA, the hexamer AAAAAA recurs 4 times, capturing runs of nucleotides. Consequently, in genomic data, runs of nucleotides shorter than 6 bp are discarded. Similarly, in the sequence <u>ATATAT</u>ATATA the dinucleotide tandem repeats are captured by the hexamers ATATAT and TATATA, each recurring 3 times. Because motifs need not recur in tandem and all short motifs over the alphabet are evaluated, all types of repeats are considered when evaluating CR, including highly diverged repeats and long motifs. Note that many short motifs (i.e., triplets and hexamers) will recur in sequences composed of longer repeats, and hence, long repeats are also captured. Obviously, the longest motif that can be identified is limited by half the short read length. For further details on how the developed procedure captures all types of repeats, from runs of amino-acids to long repetitive domains, in protein sequences, see (Persi & Horn, 2013). Examples of protein repeats and their corresponding coding DNA repeats, in repeat unstable genes identified in this study, are provided in **Figures S11-S12**.

Importantly, other genomic aberrations, such as segmental duplication and gene copy-number alterations, can alter CR if they encompass compositionally ordered regions (in which motifs are recurrent within the short-read length). Therefore, CR should be interpreted as a global measure of the repeat content that weighs in the contribution of other somatic aberrations that involve repeat-containing proteins. However, in this study, we have found that repeat-instability was independent of copy-number alterations or of ploidy (cf. Figures 3 and 5). Hence, in practice, CR is weakly sensitive to such changes in the analyzed datasets.

### Application of CR Analysis to Proteomic Datasets

Proteomic data of tumor samples and their matched-normal samples were obtained from 21 breast cancer patients (Table 1). In 10 patients, both stage III and metastatic sites were obtained, with 52 samples in total: 21 matched-normal, 11 stage II primary tumor, 10 stage III primary tumor and 10 metastatic. Mass spectroscopy SILAC-based measurements were performed in 2 different experimental pools, of 11 and 10 patients respectively, where each pool included patients with metastatic spread. The proteomic raw data for these two pools included ~120k and ~140 identified peptides, respectively. Peptide lengths typically vary between 10-30 amino-acids. We estimated the CR of each triplet *m* in each sample based on the list of identified peptides (*i* = *1*…*Ns*), taking the normalized SILAC intensity ratio (i.e., relative to a reference sample) of peptide *i* as the weight, *w_i_*, in equation 1.

### Application of CR Analysis to Genomic Datasets

To analyze the genomic datasets (Table 1), CR evaluation was applied directly to Fasta-q files of each sample, generated from BAM files using Samtool and Bedtool. In all datasets, the length of the DNA short reads was 76 base-pairs. We assumed no prior knowledge of the directionality such that the entire raw data, including unmapped reads, was considered for CR evaluation. In each short-read, a nucleotide that had a probability of error *P_err_* > 0.05, as specified by its Phred-score, was labeled as *‘N’*. Then, a search for valid DNA *k*-mers (i.e., *k*-mers that do not contain *N*) was performed. The effective number of invalid short-reads in a sample, *E*, that is, the number of base-pairs with *P_err_* > *5%* divided by 76, the short-read length, was of the order of 5-10% in these datasets. The length of short reads sets an upper bound on the possible number of recurrences within a short-read, and in turn on CR. Hence, for comparison across studies, it is essential that the short read length of NGS is identical. Further, we found that different sequencing kits affect CR evaluation. Thus, we performed all our analysis based on samples that where sequenced with the same kit (SureSelect).

In all datasets, coverage depth was approximately *x50-100*. With such large coverage depth, CR is highly stable, as demonstrated by its saturation at very low coverage depths, and the fast convergence of the statistical errors of CR to a narrow distribution, as more and more data (i.e., short-reads) is considered for CR evaluation (**Figure S2**). These statistical errors may arise due to various effects, such as unequal sampling of genomic regions. Nonetheless, empirical estimation of the statistical error of CR shows that it is relatively small (mean and median ~ 10^-4^), and consequently, by propagation of errors, the statistical error of the repeat instability signature (ΔCR) is also of the order 10^-4^ (that is, 0.01%). Note that a somatic change in the DNA (e.g., an expansion of a motif) will be sampled proportionally to the coverage depth and will therefore be identified, over the background of the statistical fluctuations.

### Benign Prostate Hyperplasia Dataset

After informed consent, benign prostatic hyperplasia (BPH) tissue samples were collected from patients who underwent surgical resection due to symptomatic BPH. Samples were examined and annotated by expert genitourinary pathologists. Tissue cores were taken for DNA extraction followed by whole exome or whole genome sequencing (>40X). Matched blood samples were used as patients’ control samples. The study was approved by the Weill Cornell Medicine IRB. A manuscript on BPH comprehensive molecular characterization including this study data is in preparation [Liu D et al., in preperation].

### Mapping Motifs to Genes and Estimating Repeat-Instability in Genes

To map motifs to genes, we extracted from the raw data those sequences in which motifs are highly recurrent (≥2 in peptides, ≥4 in DNA short reads), and mapped them to the human genome/proteome. Evaluation of CR for genes is not practical because, unlike motifs, only a small fraction of the reads map to a given gene, resulting in significant noise in the estimates of CR at the gene level. Alternatively, we calculated the total number of recurrences of all motifs across all sequences mapped to a gene in tumor samples (*A_tumor_*) and in a reference tissues, either adjacent matched-normal tissue or blood (*A_Ref_*). Then, we defined the *repeat-instability* of a gene by Δ*A* = *A_tumor_ - A_Ref_*, in a patient. Δ*A* signifies the difference in the actual number of recurrences of motifs in the sequences belonging to a gene, between the tumor and the reference tissue. To allow for a comparison between samples and patients, *A*, was always normalized by the number or total reads, as explained next.

In genomic data, we mapped all compositionally ordered sequences to the human genome using BWA, assigning to each read the most likely gene to which it belongs. Then, for each gene, we estimated the total weighted recurrence in each sample as: *A = Ʃ_h_ WR_h_*, where *R_h_* is the total number of recurrences of hexamer *h* in all the mapped sequences, and *W* = *1/(N_sr_-E*) is a constant weight factor of the relative abundance of short-reads in a sample: *N_sr_* is the total number of short-reads in the entire sample, and *E* is the number of invalid short-reads, such that *A* is comparable between two samples of different size and across patients. The mapping of two sets of hexamers, the 154 discriminative hexamers and dominant hexamers (cf. Table 2 and Figure 3) is shown in **Figure S9**. Then, we assessed Δ*A* for all genes containing discriminative hexamers. Genes that were identified as repeat-unstable in at least 75% of the patients in the Barbieri dataset (n=3321) were considered for enrichment analysis with known cancer genes (**Figure S10**).

In proteomic data, mapping triplets is straightforward because the vast majority of triplets occur in only one or a few peptides, and each peptide is maps uniquely to one protein. This mapping is provided by the output of MaxQuant which compares the similarity of peptides to proteins in the UniProt database. *A* is weighted by the measured abundance intensities of each peptide, such that *A = Ʃ_t_ W_t_R_t_*, where *W_t_* is the intensity of the peptides matched to a gene in which the triplet *t* recurs *R_t_* times. Two maps for each experimental pool (described above) were built (for a set of discriminative triplets). We considered a valid mapping (triplets to peptides to genes) only if in both experimental pools a given triplet recurred in identical peptides (and thus genes). We mapped the discriminative triplets identified by Fisher-score and Kolmogorov-Smirnov tests, by CR-based criteria, or by their association with cancer progression (Figure 2, and **Table S1**). For enrichment analysis, we selected those triplets that have consistent variation across patients (i.e., CR-increase in >70% of the patients or CR-decrease in >70% of the patients, as in Figure 3B). These triplets mapped to 491 proteins which are enriched with 63 known cancer genes (**Figure S6**).

### Statistical and Machine Learning Tools

Principal component analysis (PCA) was applied to the matrix of CR evaluations, of patients (observations) *x* motifs (features). The CR matrix was standardized such that each column is normalized by its standard deviation across observations. Support vector machine (SVM) with linear Kernel function was used for the classification tasks in a standard leave-one out procedure.

## Supporting information

## Authors Contributions

EP, FD and DH conceived and designed the study. EP and DH developed the methodology. EP and DP performed the analysis with the assistance of YIW. All authors participated in the design of the study and interpretation of the results. YP and TG contributed the proteomic dataset. DP, CB, PG, BMF, HB, MAR and FD contributed the genomic datasets. EP, EVK, FD and DH wrote the paper with the assistance of DP, YIW, TG and contributions from all authors. EP, FD and DH co-directed the study.

## Acknowledgments

We thank Eytan Ruppin, members of the CBCB at UMD, members of the Koonin group at the NIH for insightful discussions and feedback. The study was partially supported by intramural funds of the US Department of Health and Human Services (to the National Library of Medicine) and by the Centre for Integrative Biology of the University of Trento. EP was partially supported by the Safra Center for Bioinformatics at Tel Aviv University and by the Terrapin project at UMD.

