## Supplementary material for "Proteomic and Genomic Signatures of Repeat-instability in Cancer and Adjacent Normal Tissues"

#### Supplemental Figures

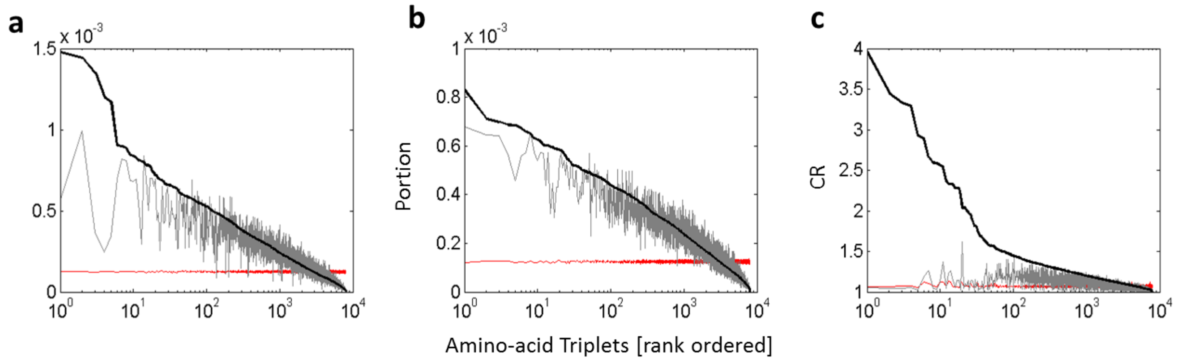


**Figure S1**: **Significance of the CR measure.** Evaluation of three measures of triplet recurrences on the human proteome (black) and on two random models, uniform (red) and human-unigram (grey) are shown in each subfigure. **A)** the normalized number of total recurrences of each triplet, i.e. the triplet frequency. **B)** the normalized number of sequences on which each triplet appeared, i.e. the portion of the sequences in the proteome on which a triplet recurs. **C)** the compositional order ratio (CR) as defined in equation 1. In all subfigures triplets are ordered by their respective measure. Only CR allows for a robust identification of repetitive triplets that are not expected at random.

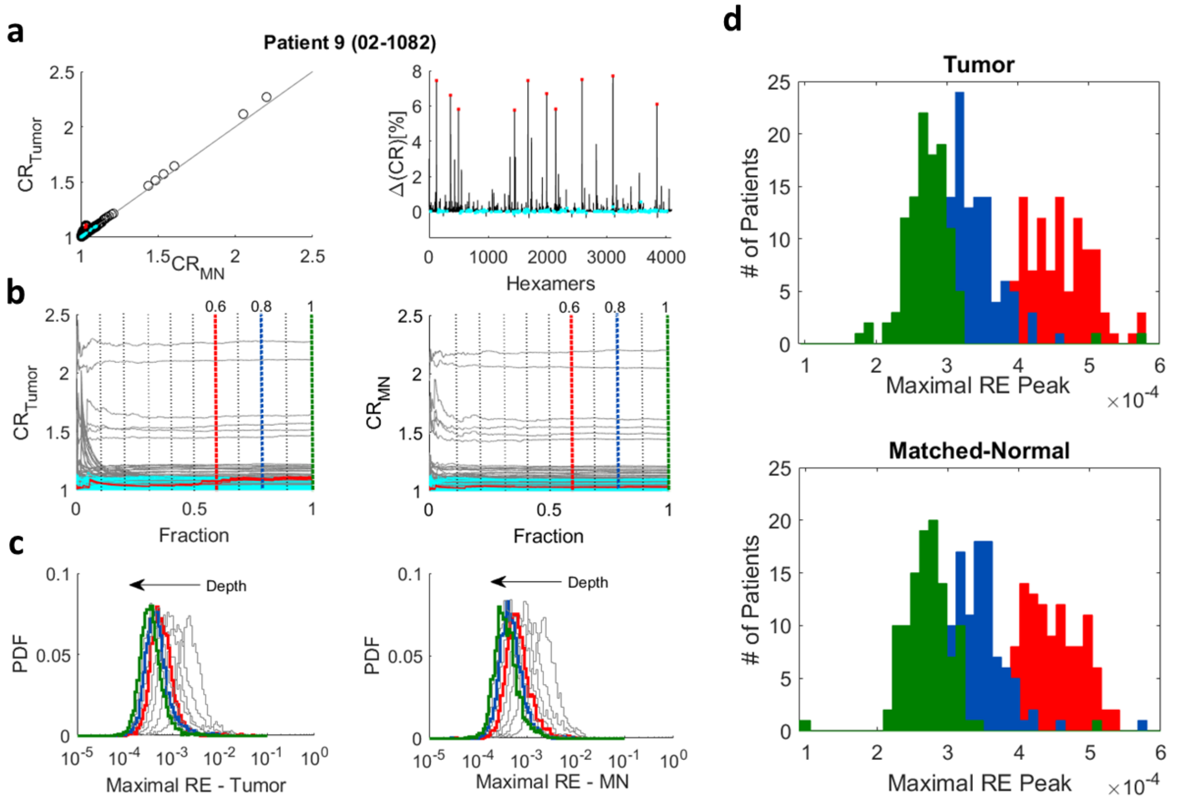


**Figure S2**: **Stability and significance of the CR signal in a single patient**. **A)** Analysis of one prostate cancer patient (ID: 02-1082) from Barbieri et al. (2012). CR of hexamers of the tumor sample vs. the matched-normal (MN) adjacent tissue sample show high correlation, as expected at the DNA level (*left*). The respective **repeat instability** signature of tumor vs. normal is shown on the right. Dominant hexamers are marked in red and discriminative hexamers are marked in cyan (see **Table 2** and **Figure 3**). **B)** CR of hexamers as a function of coverage depth. Coverage depth is shown as the fraction ($F$ = 0-1) of the short-read data considered for CR evaluation in the tumor sample (*left*) and in the MN sample (*right*). In this patient, the total number of short-reads is 267,530,730 (F=1) in the tumor sample and 275,879,972 (F=1) in the MN sample. X-axis is binned, such that each point corresponds to an accumulation of 375,000 short-reads. Dominant and discriminative hexamers are colored in red and cyan, respectively. The order of short reads was randomized before evaluating CR. Note that **CR reaches a plateau** as more data is considered, indicating that CR is a stable measure. **C)** Distribution of the **maximal relative error**, defined as |CRmax – CRmin|/CRmin evaluated from the fluctuations of CR in the interval [$F$-0.1, $F$], for each depth $F$. Distributions for depths $F$=0.6 (blue), 0.8 (blue) and 1 (green) are highlighted. **D)** The peak of the distribution of the maximal relative error was then inferred for each of the samples in the Barbieri et al (2012) cohort. The distribution of these peaks for the 111 tumors (*top*) and 111 matched-normal samples (*bottom*) are shown for the 3 depths: $F$=0.6, 0.8 and 1. Note that because CR is of the order of ~1, the relative error is proportional to the absolute error. Hence, by propagation of errors, the statistical error of ΔCR is at range of 0.01%-0.1%.

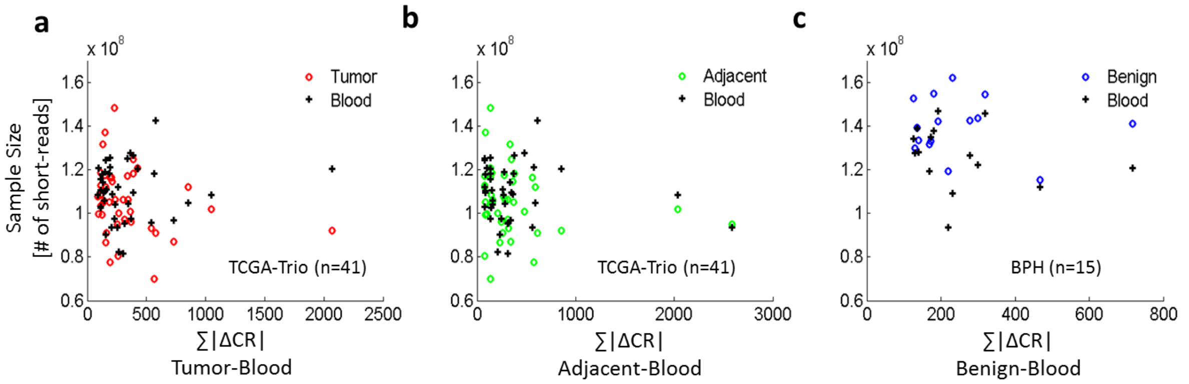


**Figure S3**: **Amplitude of repeat-instability signatures as function of the sample size**. The amplitude of signatures measured as ∑|ΔCR| (i.e., the area under the signature curve summed over all hexamers) is shown against the sample size (in total number of short reads) for each patient and tissue, using the TCGA prostate dataset and the benign hyperplasia dataset. **A)** ∑|ΔCR| of tumor vs. blood signatures in the TCGA database. **B)** ∑|ΔCR| of tumor vs. adjacent (MN) signatures in TCGA database. **C)** ∑|ΔCR| of benign vs. blood signatures. There is no dependence between ∑|ΔCR| and the sample sizes, testifying that the overall signature is unbiased with respect to the sample size (i.e., sequencing depth). This independence was observed in all datasets.

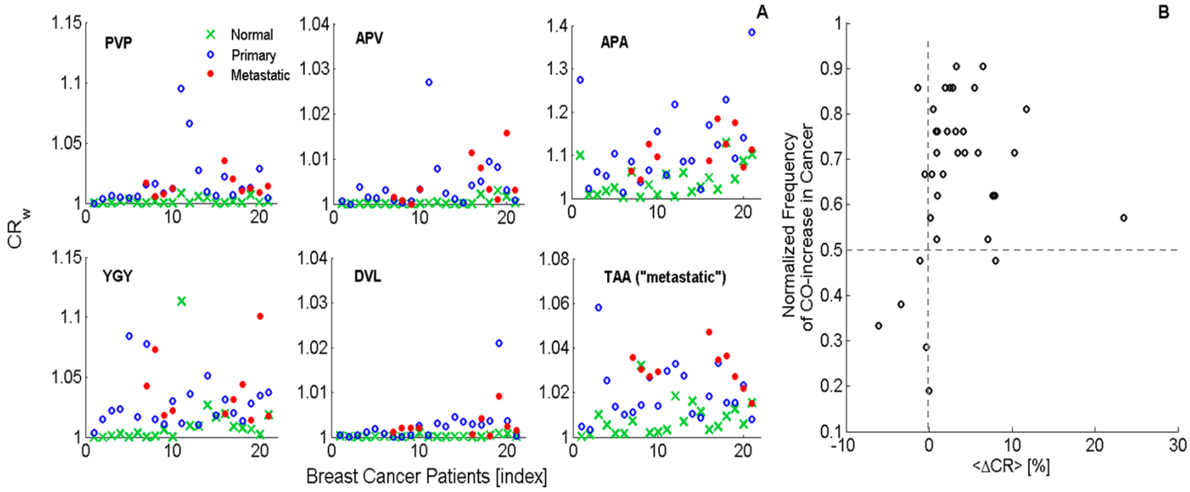


**Figure S4**: **Examples of highly discriminative triplets in the breast cancer proteomic dataset**. **A)** The CR of the triplets `PVP`, `APV`, `APA`, `YGY`, `DVL`, `TAA`, across the 21 breast cancer patients, showing clear changes from the matched-normal (green) tissues to cancerous tissues (blue: primary stage II-III; red: metastatic). **B)** The normalized frequency of CR-increase vs. the average change in CR across patients (<ΔCR>), for the 36 selected high CR triplets. Note that these features appear mostly in the second and third quarters; hence, triplets with high CR are significantly and frequently altered in cancer.

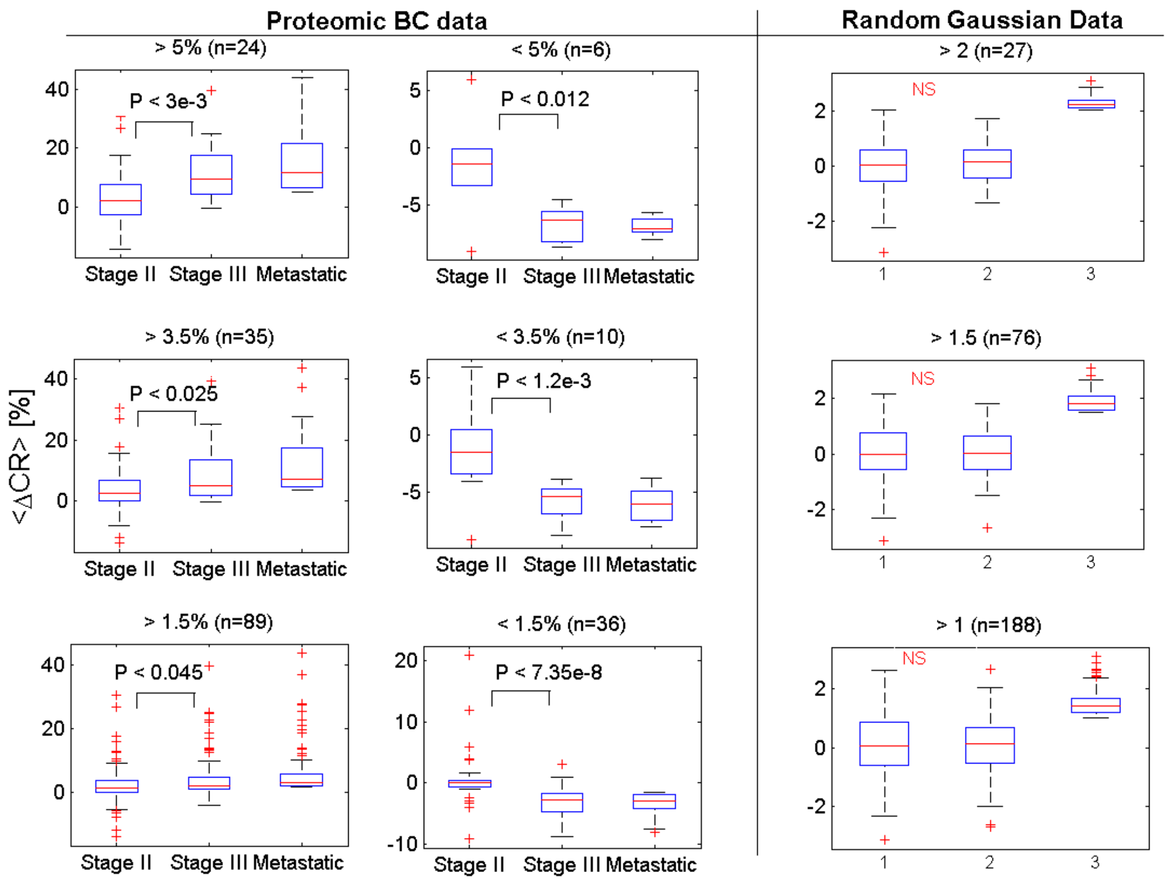


**Figure S5. Correlation between repeat-instability signatures and tumor progression in the proteomic breast cancer dataset**. **Left)** In the proteomic data, selecting for triplets that increase (first column) or decrease (second column) in the metastatic samples, always results in an evident progression from stage II to stage III (to metastatic). **Right)** to ensure that this progression is real we validated on a random matrix of similar dimension that selecting of a given group (3) never results in seen difference between that other two groups (1 & 2). P-values correspond to Kolmogorov-Smirnov test for rejecting the null hypothesis that the two distributions are drawn from a similar underlying distribution. From bottom to upper panels, plots are shown for increasing thresholds on |ΔCR| used for selecting triplets that increase/decrease in the metastatic (and simulated data). NS = not significant.

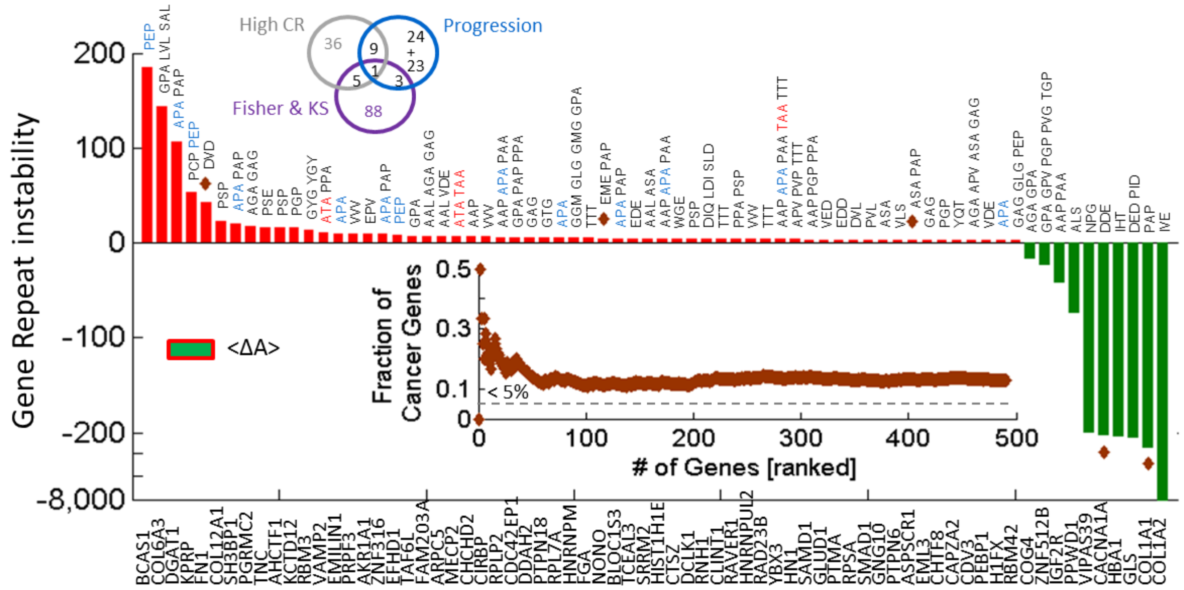


**Figure S6**: **Enrichment of repeat unstable genes containing discriminative triplets in the proteomic breast cancer dataset**. We mapped peptides containing discriminative triplets (van diagram, and see also **Table S1**) to proteins in the proteomic dataset of breast cancer (**Methods**). The analysis identified 491 repeat-unstable genes, out of which 63 are known cancer genes (HG *P*-value < 6e-15). The top 70 genes ranked by mean repeat-instability across patients (<Δ*A*>) are shown. Cancer genes (n=903) are marked by brown diamonds. Inset shows the fraction of cancer genes as function of the number of repeat-unstable genes (ranked by <|Δ*A*|>) showing enrichment relative to the background (<5% of human proteome). The most repeat-unstable gene is the breast carcinoma amplified sequence, *BCAS1*, through the recurrence of the triplet PEP which is associated with cancer progression. Within the top 100 repeat-unstable genes we identify *FN1*, *NONO*, *ASPSCR1*, *RBFOX2*, *FUS*, *COL2A1*, *VIM*, *MYD88*, *PSMA1*, *RPLP0*, *SMCHD1*. Further, other identified genes include the gene *TNC* which plays a crucial role in metastasizing from the breast to the lung. The triplet APA, considered discriminative by all criteria, points to *SH3BP1* which binds preferentially to the *ABL1* proto-oncogene. APA also points to the ribosomal protein RPL7A which is involved in the activity of tyrosine kinase (*trk*) proto-oncogene. The metastatic-specific triplets TAA and ATA (see **Figure S4**) point to *MECP2*, the oncogene *RBFOX2* which regulates estrogen receptor 1 transcriptional activity, *HLA-B* which is differentially expressed in tumors and involved in immunity escape, necessary for cancer progression, and *STAT3*.

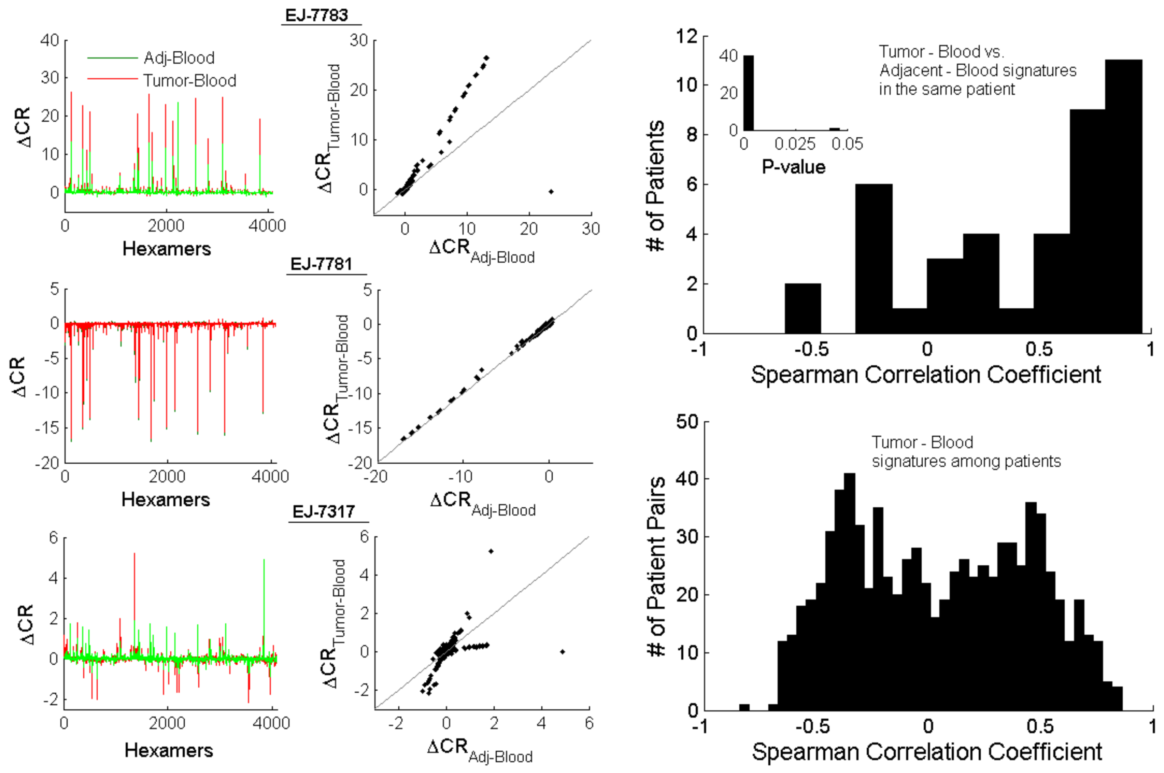


**Figure S7. Analysis of the signature of prostate samples (tumor and matched-normal adjacent tissues) in the TCGA dataset measured relative to the blood sample from the same patient. Left)** Examples of tumor tissue signatures (red) and matched-normal (MN) adjacent tissue signatures (green), measured relative to the blood, in 3 typical patients. The correlation between the signatures is also shown (middle panel). Note that in a given patient the signatures are usually either mostly positive or mostly negative, representing an increase or decrease of the CR, respectively, however, regardless of this directionality the correlation is positive in most cases. The latter indicates that the signatures directionality represents an evolutionary path, of duplication or deletion of recurring motifs, and in each patient one of these two possible paths occurs. **Right)** Upper corner depicts the histogram of the Spearman correlation between tumor and adjacent normal signatures (relative to blood) within each patient. **The evident skew towards positive correlation values indicates that within the same patient the matched-normal adjacent tissue signature is quite similar to that of the respective tumor signature**. Hence, (i) the signatures are highly tissue-specific, and (ii) the direction of the signatures (i.e., dominance of expansion or contraction) is biological, otherwise, significant negative correlations would be observed due to random biases. Lower corner shows the pairwise correlation of the tumor signatures among patients, indicating similarities in the repeat instability also among patients.

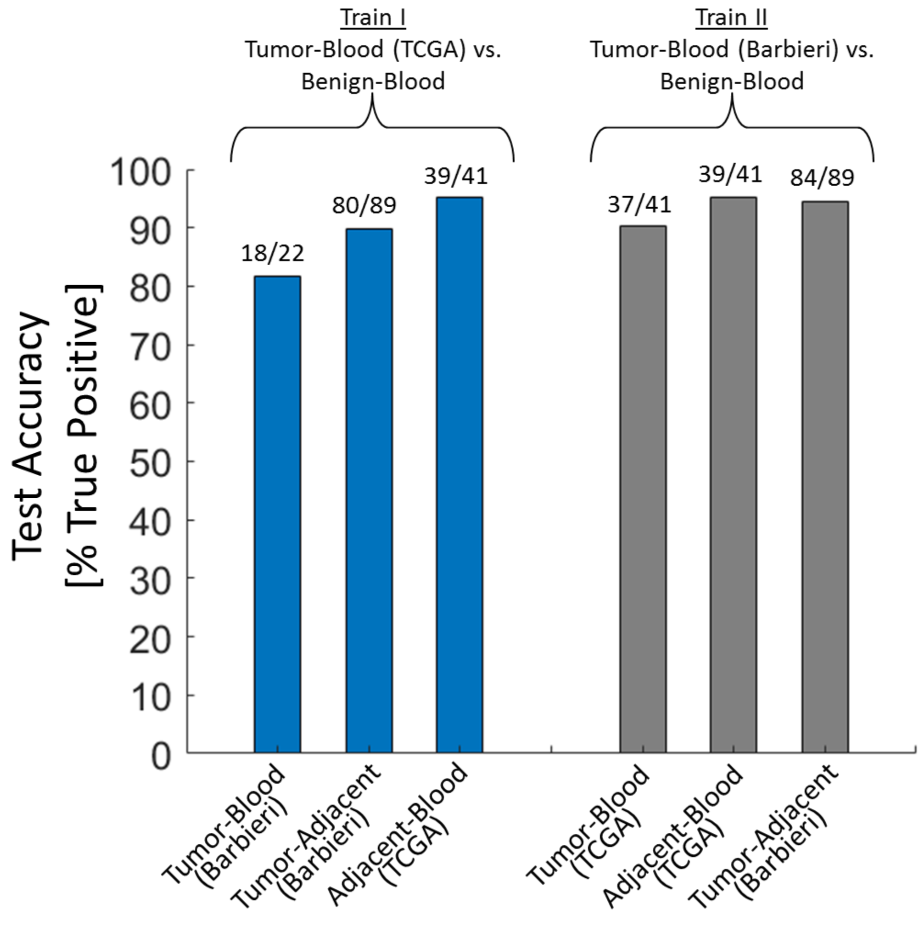


**Figure S8**: As another validation for the capacity of discriminative hexamers to predict the existence of a tumor signature, we tested the performance of SVM classifiers on 3 different signatures that we expect to contain a tumor signal based on **Figure 3** (and **Figure S7):** Tumor vs. Blood, Tumor vs. matched-normal adjacent tissue (Adjacent), and Adjacent vs. Blood. An ideal classifier would assign each of these signatures as a tumor signature (i.e., 100% true positives). Two analyses were performed. First, we trained classifiers by comparing the Tumor-Blood signature **in TCGA** (n=41) database with the Benign-Blood signature (n=15), based on the 154 discriminative hexamers identified by KS test (**Table 1**). Then, we tested the performance of the trained classifier on the three (predicted tumor) mutually exclusive signatures (indicated on the x-axis). All tests display high accuracy (*Blue bars*). Second, we trained the SVM classifier by comparing the Tumor-Blood signature **in Barbieri** database (n=22) with the Benign-Blood signature (n=15). Here we identified only 55 discriminative hexamers using KS test. Nonetheless, testing the performance of the trained classifier on the three (predicted tumor) signatures (x-axis) also led to high accuracy in identifying tumor signatures (*Grey bars*). Hence, both Tumor-Blood and Adjacent-Blood signatures display tumor characteristics, which distinguish them from Benign-Blood signatures.

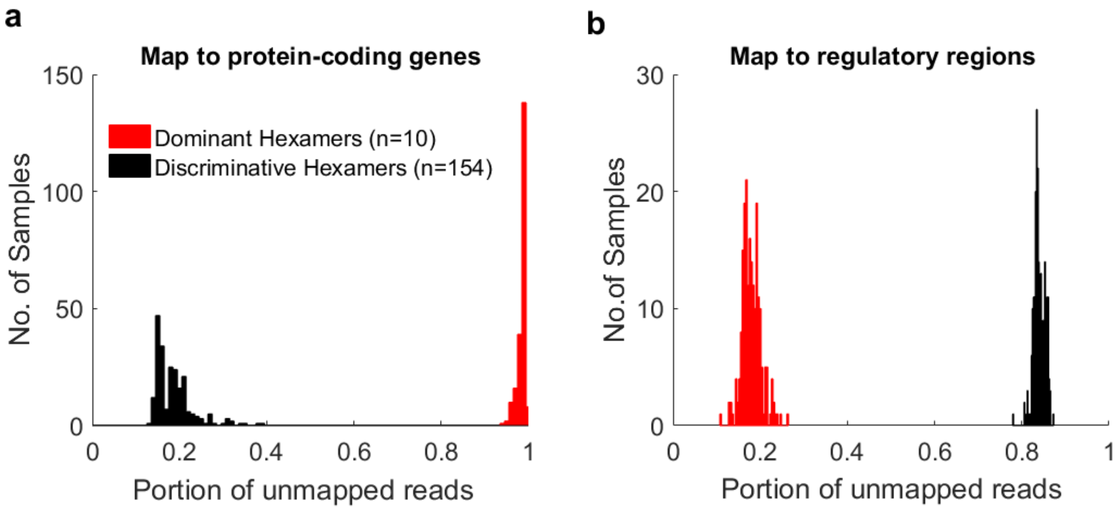


**Figure S9**: **Mapping of motifs to the human genome. A)** Mapping of identified dominant (n=10, red) and discriminative (n=154, black) hexamers in the Barbieri et al. dataset (cf. **Figure 3**) to genes **(Methods)**. Most of the reads containing at least 4 recurrences of a discriminative hexamer were mapped to protein coding genes (~20% unmapped), while most of the reads containing at least 4 recurrence of a dominant hexamers were not mapped to genes (>95% unmapped). **B)** Similar analysis was performed, this time mapping the same reads to regulatory regions. Regulatory regions were obtained from ENCODE, containing about 112K regions. Here, most of the reads containing dominant hexamers were mapped to regulatory regions (~20% unmapped), while most of the reads containing discriminative hexamers were not mapped to regulatory regions (>85% unmapped).

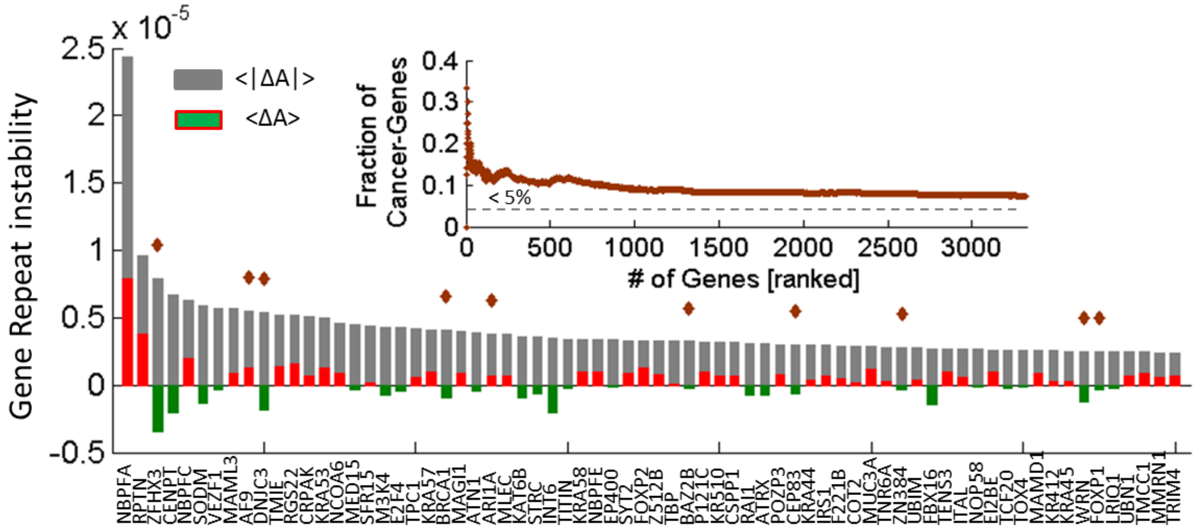


**Figure S10**: **Enrichment of repeat unstable genes containing discriminative hexamers in the Barbieri dataset**. Based on the reads that mapped to genes, we calculated the repeat instability at the gene level, $\Delta A$, in each patient (**Methods**). In the genomic prostate cancer dataset of Barbieri et al. (**Table 1**) we identified 3221 repeat-unstable genes, mapped by discriminative hexamers, with frequency >75% (i.e., repeat unstable genes in more than 75% of the patients). 246 of these genes are known cancer genes (HG *P*-value ≈ 0). Known cancer genes were taken from COSMIC and intogen (n=903; <5% of the human proteome). In the figure, the top 70 genes, ranked by the average overall repeat-instability across patients (<|$\Delta A$|>, grey) are shown. Colored bars showing the average gain or loss (<$\Delta A$>) are superimposed. Known cancer genes are marked by brown diamonds. The inset shows the fraction of known cancer genes as function of the top ranked repeat unstable genes (ranked by <|$\Delta A$|>), for all genes. Background level of cancer genes in the human proteome (<5%) is marked with a dashed line. The neuroblastoma gene *NBPFA* exhibits the largest instability. Within the top 100 most variable genes, 15-30% of the genes were known cancer genes, including: *ZFHX3*, *AF9*, *DNJC3*, *BRCA1*, *ARI1A*, *BAZ2B*, *CEP83*, *ZN384*, *WRN*, *FOXP1*, *MAML2*, *PERQ2*, *ASXL2*.

#### FOXP2(O15409)- Forkhead box protein P2

MMQESATETISNSSMNQNGMSTLSSQLDAGSRDGRSSGDTSSEVSTVELLHL**QQQQ**ALQAARQLLL**QQQ**TSGLKSPKSSDKQRPLQ
VPVSVAMMTPQVITP**QQ**M**QQ**IL**QQQ**VLSP**QQ**LQALL**QQQQ**AVML**QQQQ**LQEFYKK**QQ**EQLHLQLL**QQQQQQQQQQQQQQQQQQQQQ**
**QQQQQQQQQQQQQQQQQQQ**HPGKQAKE**QQQQQQQQQQ**LAA**QQ**LVF**QQQ**LLQM**QQ**L**QQQQ**HLLSLQRQGLISIPPGQAALPVQSLPQ
AGLSPAEI**QQ**LWKEVTGVHSMED…(282:715)

##### Discriminative hexamers

AGCAGC
 GCAGCA
  CAGCAG

##### Coding DNA

```
 M   M   Q   E   S   A   T   E   T   I   S   N   S   S   M   N   Q   N   G   M   S   T   L   S
ATG|ATG|CAG|GAA|TCT|GCG|ACA|GAG|ACA|ATA|AGC|AAC|AGT|TCA|ATG|AAT|CAA|AAT|GGA|ATG|AGC|ACT|CTA|AGC|
 S   Q   L   D   A   G   S   R   D   G   R   S   S   G   D   T   S   S   E   V   S   T   V   E
AGC|CAA|TTA|GAT|GCT|GGC|AGC|AGA|GAT|GGA|AGA|TCA|AGT|GGT|GAC|ACC|AGC|TCT|GAA|GTA|AGC|ACA|GTA|GAA|
 L   L   H   L   Q   Q   Q   Q   A   L   Q   A   A   R   Q   L   L   L   Q   Q   Q   T   S   G
CTG|CTG|CAT|CTG|CAA|CAA|CAG|CAG|GCT|CTC|CAG|GCA|GCA|AGA|CAA|CTT|CTT|TTA|CAG|CAG|CAA|ACA|AGT|GGA|
 L   K   S   P   K   S   S   D   K   Q   R   P   L   Q   V   P   V   S   V   A   M   M   T   P
TTG|AAA|TCT|CCT|AAG|AGC|AGT|GAT|AAA|CAG|AGA|CCA|CTG|CAG|GTG|CCT|GTG|TCA|GTG|GCC|ATG|ATG|ACT|CCC|
 Q   V   I   T   P   Q   Q   M   Q   Q   I   L   Q   Q   Q   V   L   S   P   Q   Q   L   Q   A
CAG|GTG|ATC|ACC|CCT|CAG|CAA|ATG|CAG|CAG|ATC|CTT|CAG|CAA|CAA|GTC|CTG|TCT|CCT|CAG|CAG|CTA|CAA|GCC|
 L   L   Q   Q   Q   Q   A   V   M   L   Q   Q   Q   Q   L   Q   E   F   Y   K   K   Q   Q   E
CTT|CTC|CAA|CAA|CAG|CAG|GCT|GTC|ATG|CTG|CAG|CAG|CAA|CAA|CTA|CAA|GAG|TTT|TAC|AAG|AAA|CAG|CAA|GAG|
 Q   L   H   L   Q   L   L   Q   Q   Q   Q   Q   Q   Q   Q   Q   Q   Q   Q   Q   Q   Q   Q   Q
CAG|TTA|CAT|CTT|CAG|CTT|TTG|CAG|CAG|CAG|CAG|CAA|CAG|CAG|CAG|CAG|CAA|CAA|CAG|CAG|CAA|CAA|CAG|CAG|
 Q   Q   Q   Q   Q   Q   Q   Q   Q   Q   Q   Q   Q   Q   Q   Q   Q   Q   Q   Q   Q   Q   Q   H
CAG|CAA|CAA|CAA|CAA|CAA|CAG|CAG|CAA|CAA|CAG|CAG|CAG|CAG|CAG|CAA|CAG|CAG|CAG|CAG|CAG|CAA|CAG|CAT|
 P   G   K   Q   A   K   E   Q   Q   Q   Q   Q   Q   Q   Q   Q   Q   L   A   A   Q   Q   L   V
CCT|GGA|AAG|CAA|GCG|AAA|GAG|CAG|CAG|CAG|CAG|CAG|CAG|CAG|CAA|CAG|CAA|TTG|GCA|GCC|CAG|CAG|CTT|GTC|
 F   Q   Q   Q   L   L   Q   M   Q   Q   L   Q   Q   Q   Q   H   L   L   S   L   Q   R   Q   G
TTC|CAG|CAG|CAG|CTT|CTC|CAG|ATG|CAA|CAA|CTC|CAG|CAG|CAG|CAG|CAT|CTG|CTC|AGC|CTT|CAG|CGT|CAG|GGA|
 L   I   S   I   P   P   G   Q   A   A   L   P   V   Q   S   L   P   Q   A   G   L   S   P   A
CTC|ATC|TCC|ATT|CCA|CCT|GGC|CAG|GCA|GCA|CTT|CCT|GTC|CAA|TCG|CTG|CCT|CAA|GCT|GGC|TTA|AGT|CCT|GCT|
 E   I   Q   Q   L   W   K   E   V   T   G   V   H   S   M   E   D   . . . (281:715)

GAG|ATT|CAG|CAG|TTA|TGG|AAA|GAA|GTG|ACT|GGA|GTT|CAC|AGT|ATG|GAA|GAC| . . . (843: 2145)
```

**Figure S11**: **Example of repeat unstable protein FOXP2 containing amino-acid runs**. FOXP2 was identified as a repeat unstable gene in most patients in the Barbieri dataset (cf. **Figure S10**, ranked number 35). We extracted its canonical amino-acid sequence and its respective protein-coding DNA. The first 281 amino-acids, which contain tracks of glutamine (Q) are shown, where Q is bold. The rest of the protein (282:715aa) does not contain runs of Q and is not shown, for clarity of the figure. Below the protein amino-acid sequence, the 3 discriminative hexamers identified in the Barbieri dataset (cf. **Figure 3**) that recur in this protein are shown (aligned and colored). Further below, these 3 hexamers are marked on the coding DNA, showing how they capture most of the Q tracks. This example indicates how short protein repeats such as amino-acid runs can be identified using the CR methodology based on nucleotide hexamers.

#### CRIPAK (Q8N1N5)- Cysteine-rich PAK1 inhibitor

```
MHEPSL
 1   CANVECPPAHTCPCGVPACSCAHVECPPAHT 7-37
 2   CRCGVPACSHMPMWSARLLTRAHVECPPAHT 38-68
     RVHVECPPAHVPMWSAHLLTC
 3   ADVECHLLTHVPMWSARLLTCPCGVPACSHV 90-120
     PMRSARLLTRAHAECPPAHT
 4   CPCGVPACSHVPMRSARLLTRADVECPPAHT 141-171
 5   CPCGVPACSHVPTWSARLITRAHVECSPAHT 172-202
 6   CRCGVPACSHVPMWSVRLLTRADAECPPAHT 203-233
 7   CRCGVPACSHVPMWSARLLTCRCGVPACSHV 234-264
 8   CRCGVPACSHVPMWSARLLTCRCGVPACSHV 254-284
     PMWSARLLTRAHVECPPAHT
 9   CRRGVPACSRAHMECPPAHTCHCGVPACSHT 305-335
 10  CRCGVPACSHVPMWSARLLTRAYVECPPAHT 336-366
     RAHVECPPAHTCPCGVPACSHT
 11  CPCGVPACSHKALAWWFCRFPVLPAESDAVT 389-419
VHSTHGGFLIRFYVKDPFYISLHLEIT
```

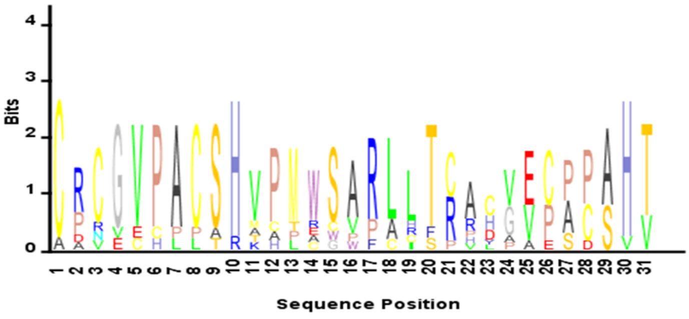


Coding DNA

```
 M   H   E   P   S   L   C   A   N   V   E   C   P   P   A   H   T   C   P   C   G   V   P   A
ATG|CAT|GAG|CCC|TCG|CTT|TGT|GCC|AAT|GTG|GAG|TGC|CCG|CCT|GCT|CAC|ACG|TGC|CCA|TGT|GGA|GTG|CCC|GCC|
 C   S   C   A   H   V   E   C   P   P   A   H   T   C   R   C   G   V   P   A   C   S   H   M
TGC|TCA|TGT|GCC|CAT|GTG|GAG|TGC|CCG|CCT|GCT|CAC|ACA|TGT|CGA|TGC|GGA|GTG|CCC|GCC|TGC|TCA|CAC|ATG|
 P   M   W   S   A   R   L   L   T   R   A   H   V   E   C   P   P   A   H   T   R   V   H   V
CCC|ATG|TGG|AGT|GCC|CGC|CTG|CTC|ACA|CGT|GCC|CAT|GTG|GAG|TGC|CCG|CCT|GCT|CAC|ACA|CGT|GTC|CAT|GTG|
 E   C   P   P   A   H   V   P   M   W   S   A   H   L   L   T   C   A   D   V   E   C   H   L
GAG|TGC|CCA|CCT|GCT|CAT|GTG|CCC|ATG|TGG|AGT|GCC|CAC|CTG|CTC|ACA|TGT|GCC|GAT|GTG|GAG|TGC|CAC|CTG|
 L   T   H   V   P   M   W   S   A   R   L   L   T   C   P   C   G   V   P   A   C   S   H   V
CTC|ACA|CAC|GTG|CCC|ATG|TGG|AGT|GCC|CGC|CTG|CTC|ACG|TGC|CCA|TGT|GGA|GTG|CCC|GCC|TGC|TCA|CAC|GTG|
 P   M   R   S   A   R   L   L   T   R   A   H   A   E   C   P   P   A   H   T   C   P   C   G
CCG|ATG|CGG|AGT|GCC|CGC|CTG|CTC|ACA|CGT|GCC|CAT|GCG|GAG|TGC|CCG|CCT|GCT|CAC|ACG|TGC|CCA|TGC|GGA|
 V   P   A   C   S   H   V   P   M   R   S   A   R   L   L   T   R   A   D   V   E   C   P   P
GTG|CCC|GCC|TGC|TCA|CAC|GTG|CCC|ATG|CGG|AGT|GCC|CGC|CTG|CTC|ACA|CGT|GCC|GAC|GTG|GAG|TGC|CCG|CCT|
 A   H   T   C   P   C   G   V   P   A   C   S   H   V   P   T   W   S   A   R   L   I   T   R
GCT|CAC|ACG|TGC|CCA|TGT|GGA|GTG|CCC|GCC|TGC|TCA|CAC|GTG|CCA|ACG|TGG|AGT|GCC|CGC|CTG|ATC|ACA|CGT|
 A   H   V   E   C   S   P   A   H   T   C   R   C   G   V   P   A   C   S   H   V   P   M   W
GCC|CAT|GTG|GAG|TGC|TCG|CCT|GCT|CAC|ACG|TGC|CGA|TGT|GGA|GTG|CCT|GCC|TGC|TCA|CAC|GTG|CCC|ATG|TGG|
 S   V   R   L   L   T   R   A   D   A   E   C   P   P   A   H   T   C   R   C   G   V   P   A
AGT|GTT|CGC|CTG|CTC|ACA|CGT|GCC|GAT|GCG|GAG|TGC|CCG|CCT|GCT|CAC|ACG|TGC|CGA|TGC|GGA|GTG|CCC|GCC|
 C   S   H   V   P   M   W   S   A   R   L   L   T   C   R   C   G   V   P   A   C   S   H   V
TGC|TCA|CAC|GTG|CCC|ATG|TGG|AGT|GCC|CGC|CTG|CTC|ACG|TGC|CGA|TGT|GGA|GTG|CCC|GCC|TGC|TCA|CAC|GTG|
 P   M   W   S   A   R   L   L   T   C   R   C   G   V   P   A   C   S   H   V   P   M   W   S
CCC|ATG|TGG|AGT|GCC|CGC|CTG|CTC|ACG|TGC|CGA|TGT|GGG|GTG|CCC|GCC|TGC|TCA|CAT|GTG|CCG|ATG|TGG|AGT|
 A   R   L   L   T   R   A   H   V   E   C   P   P   A   H   T   C   R   R   G   V   P   A   C
GCC|CGC|CTG|CTC|ACA|CGT|GCC|CAT|GTG|GAG|TGC|CCG|CCT|GCT|CAC|ACG|TGC|CGA|CGT|GGA|GTG|CCC|GCC|TGC|
 S   R   A   H   M   E   C   P   P   A   H   T   C   H   C   G   V   P   A   C   S   H   T   C
TCA|CGT|GCC|CAT|ATG|GAG|TGC|CCG|CCT|GCT|CAC|ACG|TGC|CAT|TGT|GGA|GTG|CCC|GCC|TGC|TCA|CAC|ACA|TGC|
 R   C   G   V   P   A   C   S   H   V   P   M   W   S   A   R   L   L   T   R   A   Y   V   E
CGA|TGT|GGA|GTG|CCC|GCC|TGC|TCA|CAC|GTG|CCC|ATG|TGG|AGT|GCC|CGC|CTG|CTC|ACA|CGT|GCC|TAT|GTG|GAG|
 C   P   P   A   H   T   R   A   H   V   E   C   P   P   A   H   T   C   P   C   G   V   P   A
TGC|CCG|CCT|GCT|CAC|ACA|CGT|GCC|CAT|GTG|GAG|TGC|CCG|CCT|GCT|CAC|ACG|TGC|CCA|TGT|GGA|GTG|CCT|GCC|
 C   S   H   T   C   P   C   G   V   P   A   C   S   H   K   A   L   A   W   W   F   C   R   F
TGC|TCA|CAC|ACG|TGC|CCA|TGT|GGA|GTG|CCC|GCC|TGC|TCA|CAC|AAA|GCC|CTG|GCA|TGG|TGG|TTC|TGT|AGG|TTT|
 P   V   L   P   A   E   S   D   A   V   T   V   H   S   T   H   G   G   F   L   I   R   F   Y
CCT|GTC|CTG|CCG|GCC|GAG|TCA|GAC|GCT|GTT|ACC|GTA|CAT|TCT|ACT|CAT|GGT|GGC|TTT|TTA|ATA|CGT|TTT|TAT|
 V   K   D   P   F   Y   I   S   L   H   L   E   I   T   *
GTC|AAG|GAT|CCC|TTT|TAT|ATT|TCT|CTG|CAC|CTC|GAG|ATA|ACG|TAG
```

**Figure S12**: **Example of repeat unstable protein CRIPAK containing repetitive domains**. Similarly to the analysis of FOXP2 gene above (cf. **Figure S11**), we analyzed the repeat content in the amino-acid and DNA sequences of the repeat unstable protein CRIPAK (cf. **Figure S9**, ranked number 13). This protein contains 11 repetitive domains of 31aa each (*upper*). Sequence logo is shown below. The 2 discriminative hexamers that were found on this protein (GTGCCC and GCCTGC) are marked on the protein-coding DNA (*bottom*). This example indicates how long and quite diverged protein repeats (that do not recur in perfect tandem and that diverged by accumulating mutations and indels) can be identified using the CR methodology based on hexamers.

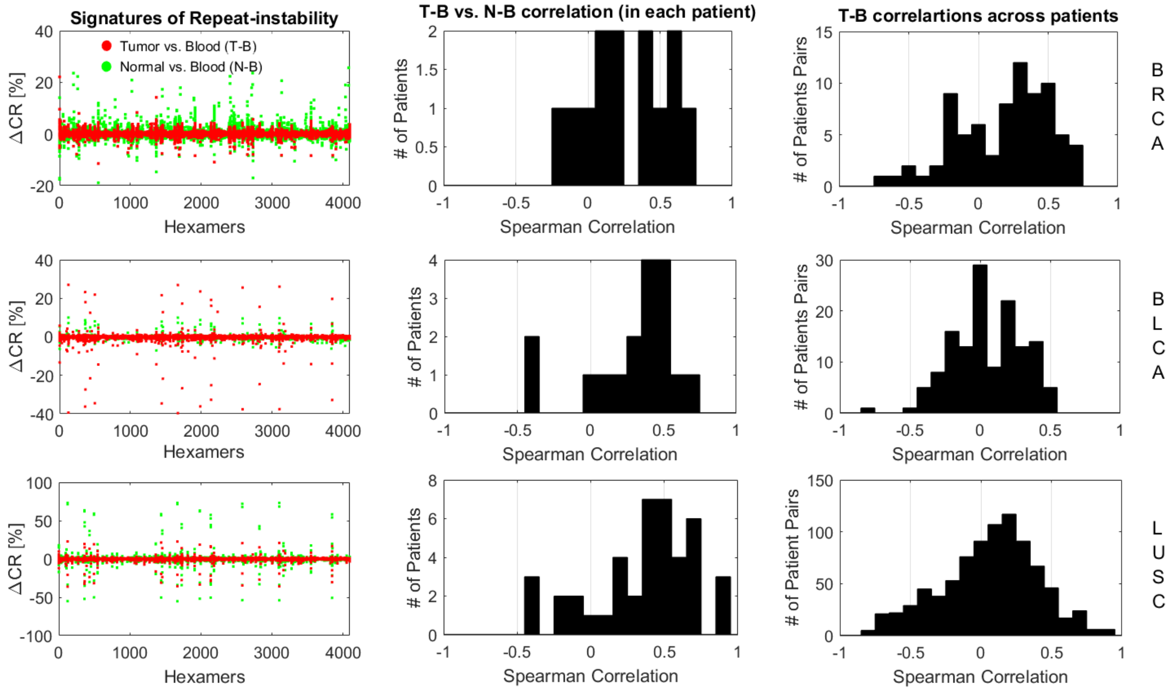


**Figure S13**: **Repeat instability signatures in tumor and corresponding adjacent matched-normal tissues in the TCGA datasets**. **Left**) The repeat instability signatures of all patients superimposed (breast, top; bladder middle; lung, bottom). **Middle**) the spearman correlation between tumor signatures (T-B) and normal signatures (N-B) measured each relative to the blood sample, in each patient. Note that correlations are positive in most of the patients, similarly to the case of prostate cancer (cf. **Figure S6**). **Right**) the pairwise correlations of the tumor signatures across patients. The bimodal distribution observed in breast and prostate cancer patients (indicating similarities across patients) weakens in the high mutation load cancers (i.e., bladder and lung).

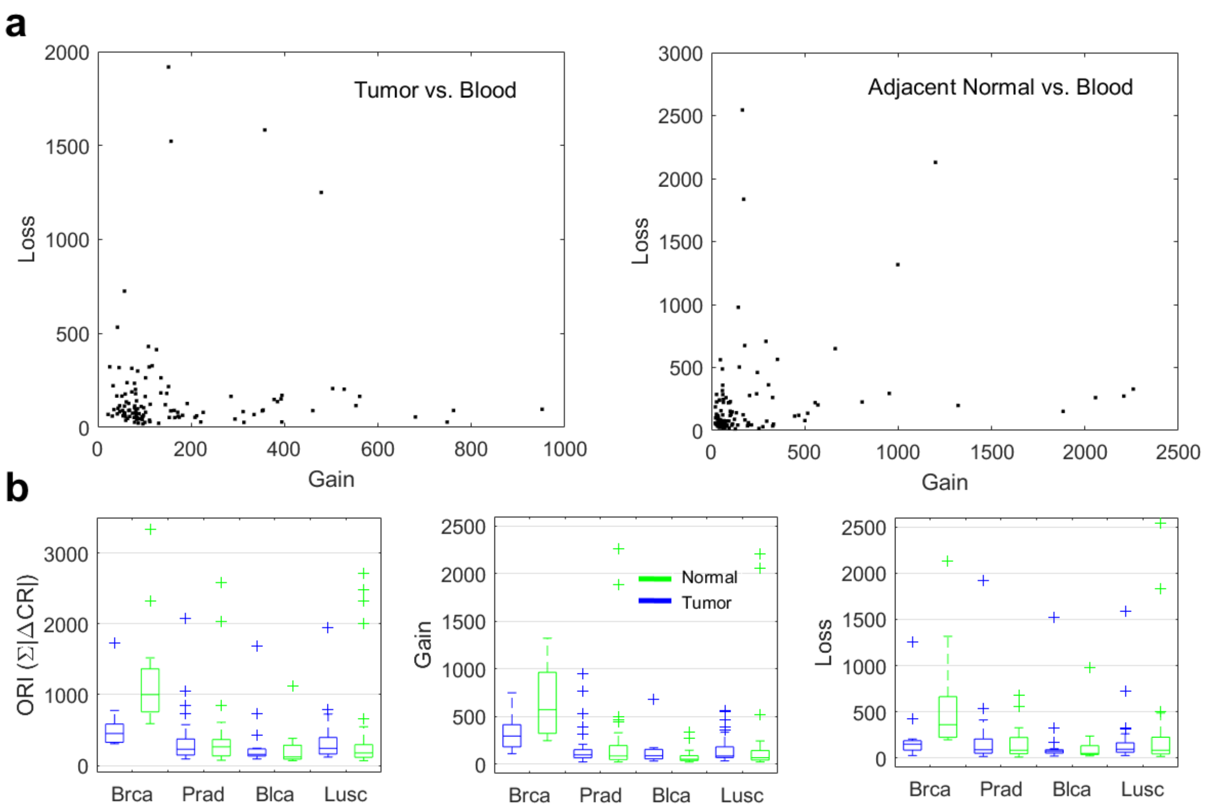


**Figure S14**: **Overall repeat instability versus gain and loss of repeats**. **A)** Scatter plots of the overall gain (Gain=∑ΔCR, for ΔCR>0) and the overall loss (Loss=∑|ΔCR|, for ΔCR<0) of repeat in tumors (*left*) and in normal tissues (*right*), for the pan-cancer TCGA data (prostate, breast, bladder and lung; **Table 1** and **Figure 4**). Tumor and normal signatures are measured relative to the blood. **B)** Distributions of the overall repeat instability (ORI=∑|ΔCR|), identical to **Figure 4** (*left*), and the overall gain (*middle*) and overall loss (*right*) in patients, across the different cancer types. On average, gain and loss are comparable, nonetheless, gain is more pronounced than loss, specifically in tumors.

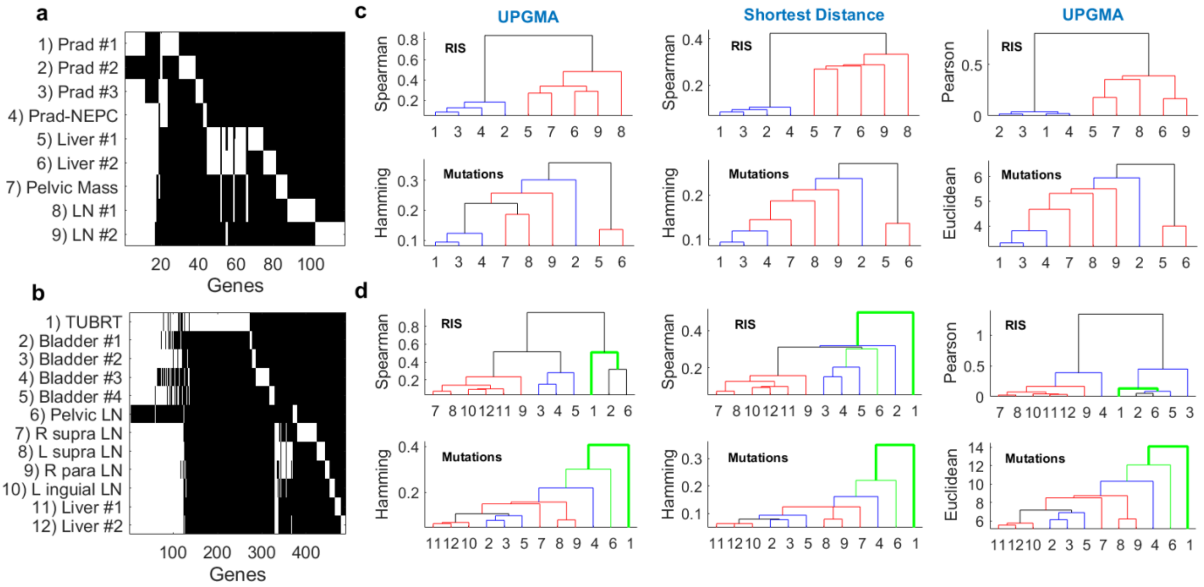


**Figure S15**: **Robustness of the distance-based phylogenies**. **A)** The non-silent mutations in the prostate cancer patient are shown across genes. Genes are ordered by those having mutations in more than one sample (shared by most), shared by primary tumors (but not metastases), private to each primary tumor, shared by metastases (but not primary) and private to each metastatic site. **B)** Distribution of non-silent mutations in the bladder cancer patient. **C)** Inferred phylogenies from the repeat instability signatures (RIS) (*top*) and from the number of non-silent point mutations (*bottom*) in the prostate cancer. The **left panel** shows the inferred phylogenies using UPGMA to link samples, as in the main text. For repeat instability phylogeny distances were calculated as 1-ρ, where ρ is the Spearman correlation. For the phylogeny inferred from the number of mutations, Hamming distances were used. The **middle panel** shows a similar analysis, using this time the shortest distance to link between samples instead of UPGMA. The results are fairly robust. The **right panel** shows the results of using UPGMA, and when the distances between samples are now calculated using the Pearson correlation (for repeat-instability inferred phylogeny) and by the Euclidean distance (for point mutation inferred phylogeny), respectively. Also here, the inferred trees are fairly robust, showing little differences from those obtained on the left panel. **D)** Similar analysis as in (C) performed for the bladder cancer patient.

### Supplemental Tables

| Feature Selection Method | Selection Criteria | # of Features | Feature Stability | Accuracy | Sensitivity | Specificity |
|---|---|---|---|---|---|---|
| none | $\Delta CR \neq 0$ | 1229 | N/A | 73% | 0.52 | 0.87 |
| Fisher-score | Top 100 | 100 | 0.83 | 81% | 0.67 | 0.9 |
| | Top 80 | 80 | 0.79 | 79% | 0.67 | 0.87 |
| | Top 60 | 60 | 0.75 | 87% | 0.76 | 0.94 |
| | Top 40 | 40 | 0.79 | 81% | 0.71 | 0.87 |
| | Top 30 | 30 | 0.73 | 79% | 0.71 | 0.84 |
| | Top 20 | 20 | 0.75 | 83% | 0.81 | 0.84 |
| | Top 10 | 10 | 0.8 | 79% | 0.67 | 0.87 |
| Kolmogorov-Smirnov | Top 100 | 100 | 0.83 | 73% | 0.62 | 0.81 |
| | Top 80 | 80 | 0.84 | 75% | 0.62 | 0.84 |
| | Top 60 | 60 | 0.85 | 71% | 0.62 | 0.77 |
| | Top 40 | 40 | 0.78 | 73% | 0.67 | 0.77 |
| | Top 30 | 30 | 0.75 | 73% | 0.67 | 0.77 |
| | Top 20 | 20 | 0.79 | 73% | 0.57 | 0.84 |
| | Top 10 | 10 | 0.77 | 77% | 0.76 | 0.77 |
| High CR | > 1.01 | 261 | 1 | 75% | 0.57 | 0.87 |
| | > 1.02 | 94 | 1 | 87% | 0.81 | 0.9 |
| | > 1.03 | 54 | 1 | 81% | 0.76 | 0.84 |
| | > 1.04 | 36 | 1 | 89% | 0.81 | 0.94 |
| | > 1.05 | 26 | 1 | 77% | 0.62 | 0.87 |
| | > 1.06 | 18 | 1 | 71% | 0.67 | 0.74 |

**Table S1**: Performance of SVM linear classifier using leave-one-out procedure, with and without feature selection. In all cases, the input to the classifier is the proteomic CR evaluations of the 52 samples obtained from 21 breast cancer patients (see Methods). The discrimination task is between the cancer samples (n=31; 21 primary tumor and additional 10 metastatic samples) and the matched-normal samples (n=21). Column I) various feature selection methods were tested. Columns II) criteria used for selecting features in each of the 52 training sets. Column III) the resulting range of number selected features in the training sets. Column IV) stability of the selected feature in the training sets, as measured by the average Jaccard score of all training set pairs (52x52). Column V) Accuracy = percentage of correct classifications. Column VI) sensitivity = TP/(TP+FN). Column VII) specificity = TN/(TN+ FP). TP = true positives, TN = true negatives, FP = false positives, FN = false negatives.
